# From aerial drone to QTL: Leveraging next-generation phenotyping to reveal the genetics of color and height in field-grown *Lactuca sativa*

**DOI:** 10.1101/2024.11.07.622452

**Authors:** Rens Dijkhuizen, Abraham L. van Eijnatten, Sarah L. Mehrem, Esther van den Bergh, Jelmer van Lieshout, Kiki Spaninks, Steven Kaandorp, Remko Offringa, Marcel Proveniers, Guido van den Ackerveken, Basten L. Snoek

**Affiliations:** Theoretical Biology and Bioinformatics, Institute of Biodynamics and Biocomplexity, Department of biology, Science Faculty, Utrecht University, Padualaan 8, 3584CH Utrecht, The Netherlands; Translational Plant Biology, Institute of Environmental Biology, Department of biology, Science Faculty, Utrecht University, Padualaan 8, 3584CH Utrecht, The Netherlands; Plant Developmental Genetics, Institute of Biology Leiden, Leiden University, Sylviusweg 72, 2333 BE Leiden, The Netherlands; Bejo Zaden B.V., Trambaan 1, 1749 CZ, Warmenhuizen, The Netherlands

**Keywords:** Lettuce, Lactuca sativa, drone, UAV, GWAS, natural variation

## Abstract

In recent years, the automation of genotyping has significantly enhanced the efficiency of genome-wide association studies. As a result, phenotyping rather than genotyping is now the rate-limiting step, especially in field experiments. For this reason, there is a strong need to further automate in-field phenotyping. Here we present a GWAS study on 194 field-grown accessions of lettuce (*Lactuca sativa*). These accessions were non-destructively phenotyped at two time points 15 days apart using an unmanned aerial vehicle. Our high throughput phenotyping approach integrates an RGB camera, a multispectral camera to measure the reflectance at 5 wavelengths (blue, green, red, red edge, near-infrared), and precise height estimation. We used the mean and other descriptives such as median, quantiles, minimum and maximum to quantify different aspects of color and height variation in lettuce from the drone images. Using this approach, we confirm several previously described QTLs, now in populations grown under field conditions, and identify several new QTLs for plant-height and color.

## Introduction

Lettuce (*Lactuca sativa L.*) is one of the most popular salad vegetables worldwide and is an excellent source of dietary minerals, fibers, and phytochemicals with related health benefits (Shi *et al*., 2022). Many different lettuce varieties are grown worldwide, and new varieties are continuously introduced (Wei *et al*., 2021; Zhang *et al*., 2017). Lettuce varieties can be grouped into several genetically and phenotypically distinguishable morphology types such as Butterhead, Cos, Latin, Cutting, Stalk, Crisp and Oilseed (Wei *et al*., 2021). Significant heritable phenotypic variation exists both within and between morphology types. For example, the leaves of different lettuce varieties exhibit a range of colors from green to red or purple (Falcioni *et al*., 2023; Su *et al*., 2020).

Precise phenotyping of color traits in field-grown lettuce, and the study of underlying genetic pathways has important implications for consumer health (Falcioni *et al*., 2022). Variation in the greenness of lettuce is partly determined by chlorophyll content. The chlorophyll content has implications for the nutritional state and quality of leafy vegetables (Heimler *et al*., 2007; Falcioni *et al*., 2023). Additionally, chlorophyll itself is considered a good source of several vitamins, important minerals, and essential fatty acids (Shi *et al*., 2022; Yang *et al*., 2022). Previously, variation in chlorophyll content in lettuce has been linked to the transcription factor LsGLK (Zhang *et al*., 2022). Additionally, anthocyanins are responsible for the red or purple coloration displayed by many lettuce varieties. Anthocyanins and other flavonoids are strongly correlated with antioxidant activity. Red lettuce varieties are rich in these flavonoids and other bioactive compounds associated with health benefits (Mampholo *et al*., 2016; Cheng *et al*., 2014). Several genes such as *Red Lettuce Leaves* 1 to 4 (*RLL1, RLL2*, *RLL3*, *RLL4*) and anthocyanidin synthase (*ANS*) have been implicated in lettuce anthocyanin content (Zhang *et al*., 2017; Su *et al*., 2020). Between-genotype variation in metabolites like chlorophyll and anthocyanin is approximated from imaging data using color intensity and vegetation indices, usually the mean intensity is used (Huang et al., 2021; Kureel et al., 2022; Falcioni et al., 2023). However, mean or median color traits are insensitive to spatial variation. For example, the mean red intensity of a green leaf with some red speckles and a homogenous slightly red leaf will be the same. To capture this spatial heterogeneity, we also use extended descriptives of color intensities and vegetation indices such as quantiles, minimum and maximum.

Another commercially important trait for lettuce is bolting time. Bolting is the transition from the vegetative growth stage to the reproductive growth stage, marked by rapid elongation of the stem (Chen *et al*., 2019). Because bolting makes the plant taste bitter, lettuce is typically harvested in its vegetative growth state. Therefore, delayed bolting and a stable flowering time are desirable traits in lettuce (Leijten *et al*., 2018). Bolting and flowering time are complex traits that have been extensively researched. Both are influenced by ambient temperature, photoperiod, the gibberellic acid pathway, age, and carbohydrate status (Han, Truco, *et al*., 2021). A total of 67 non-overlapping QTLs have been reported for bolting and flowering time (Han, Truco, *et al*., 2021). Bolting time in lettuce is strongly affected by a lettuce homolog of the Arabidopsis gene *Phytochrome C*(Rosental *et al*., 2021).

Phenotypic variation can be associated with potential underlying genomic loci using a statistical procedure called genome wide-association study (GWAS). Genetic loci associated with trait variation are referred to as quantitative trait loci (QTLs). Due to the advancement of next-generation sequencing, the bottleneck for GWAS is no longer genotyping but phenotyping (Spindel *et al*., 2018; Watanabe *et al*., 2017; Xiao *et al*., 2022). Recently, several studies have increased phenotyping efficiency by applying modern drone and sensor technology (Spindel *et al*., 2018; Han, Wong, *et al*., 2021; Ye *et al*., 2023; Ding *et al*., 2023). Drone-assisted measurements provide an opportunity for large-scale in-field phenotyping.

In this study, we performed GWAS on plant-color and height traits of 194 *L. sativa* varieties grown in a large field experiment. The traits were obtained using an aerial drone, which imaged the field at two time points 15 days apart using an RGB and a multispectral (MSP) camera and a 2.5D approach to determine plant height. The genotypes of the lettuce varieties were obtained using the assemblies from Wei *et al*., (2021) combined with 67 newly sequenced lines (Van Workum et al., 2024a). We show that the measured traits form coherent clusters with a shared genetic basis. Our approach is validated by detecting previously detected QTLs. We recover the loci for PhyC, RLL2 and ANS which were previously already detected in field conditions from Wei *et al*., (2021). Furthermore, we report novel QTLs and candidate genes affecting variation in color and height, demonstrating the power of drone-assisted phenotyping. We also show the added value of constructing many traits as ratios from the imaging data and describing phenotypes using extended descriptives beyond the mean value. QTLs can depend on the growth stage of an organism (Bac-Molenaar *et al*., 2016; Vasseur *et al*., 2014; van Eijnatten *et al*., 2024). By phenotyping twice 15 days apart we show that traits from both days uncover unique QTLs. Taken together, this study shows that high-throughput, non-destructive, drone-assisted phenotyping can be used to show the interaction between genes, environment, and development under field conditions.

## Methods

### Fields measurements

For this experiment 194 *Lactuca sativa* accessions (**Supplementary Table 1**) were grown under field conditions near Maasbree in the Netherlands. Each accession belongs to a morphology group. These groups are Butterhead, Cos, Crisp, Cutting, Latin, Oilseed and Stalk. We show an example for each group in Figure 1. The accessions shown here are LK061, LK147, LK087, LK183, LK108, LK198 and LK194 respectively. Each accession was grown on two plots, where each plot contained 30 to 40 individual plants. The plants were sowed at a commercial plant nursery on the 25^th^ and 26^th^ of March 2021 and planted in the field on the 27^th^ and 28^th^ of April 2021. The entire field was imaged on two timepoints, on the 11^th^ of June (day-78), and on the 25^th^ of June (day-93). This was done by an unmanned aerial vehicle (UAV), the DJJ Matrice 210 RTK V2, which was equipped with a conventional RGB camera and a multispectral (MSP) camera, the Micasense Altum high resolution. The MSP camera captures blue (475nm, 32nm bandwidth), green (560nm, 27nm bandwidth), red (668nm, 14nm bandwidth), red-edge (717nm, 12nm bandwidth) and near-infrared (NIR 842nm, 57nm bandwidth) light (AgEagle, 2020). Because some plants were removed between day-78 and day-93 for use in destructive measurements (not included in this study), we only examine the average height and color of the plants rather than also looking at the total surface area. The images captured by the drone were stitched together by the DroneWerkers company with geotiff software as well as the estimation of the height by 2.5D software using the overlap and difference in angle between the individual images (DroneWerkers, 2022).

**Figure 1:**
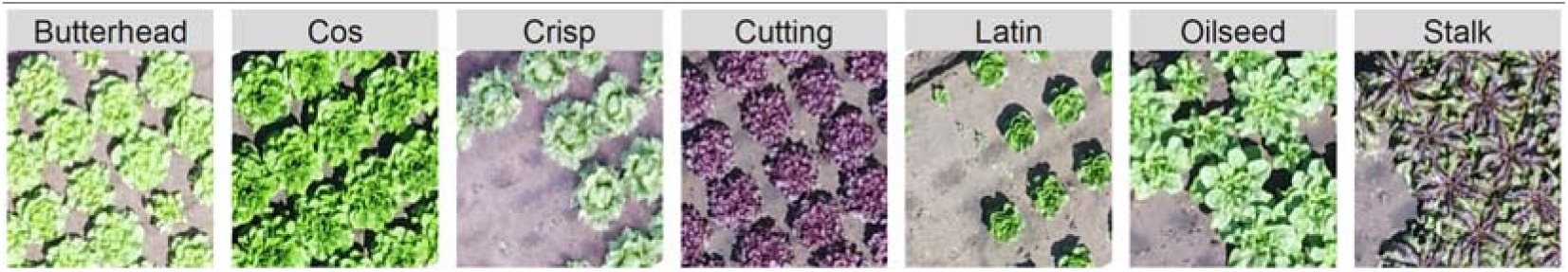
Seven different types of lettuce from the perspective of the unmanned aerial vehicle. One accession per lettuce type is shown. The name of the lettuce type is shown on top of each panel.

### Plot detection

Using the recorded geographic coordinates, individual plots within the field were isolated from the full image. Pixels in the image corresponding to plants were identified by a thresholding approach using the enhanced vegetation index (EVI). We used different EVI thresholds on the two imaging days, 0.25 and 0.4 respectively, due to differences in the overall brightness of the images between days. These thresholds were visually determined to represent the best tradeoff between not capturing soil pixels while still correctly identifying most plant pixels (**Supplementary Figure 1).** The Enhanced Vegetation Index (EVI) more effectively identified plant pixels compared to the NDVI or SR, because it was better at distinguishing between the dark red/purple accessions and soil pixels covered in shadow.

### Trait extraction

After isolating the plant pixels per plot we calculated the color- and height trait traits per plot (30 to 40 plants). For each trait, we characterized the distribution per plot using the mean but also other descriptives such as the trimmed means (excluding either the top and bottom 5%, 10% or 40%) the median, the 5%, 10%, 25%, 50%, 75%, 90% and 95% quantile, the minimum and maximum values, and the skewness and kurtosis of the distribution (**Supplementary Table 3**). We used the RGB and MSP values directly as traits for GWAS. Additionally, we used the MSP data to construct several vegetation indices such as the normalized difference vegetation index (NDVI), the structure insensitive pigment index (SIPI), the atmospherically resistant vegetation index (ARVI), the chlorophyll index red-edge (CIred), the simple ratio (SR), the enhanced vegetation index (EVI), the normalized difference red edge index (NDRE) and the weighted difference vegetation index (WDVI) (**Table 1**). We also calculated different ratios of the RGB values, by taking 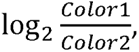 and used these as phenotypes (**Supplementary Table 2**). We quantified the height by subtracting the mean elevation of the soil pixels of each plot from the height of each plant pixel to account for the unevenness of the terrain. To calculate the final trait values, we took the mean of the replicate plots per trait. All these traits were calculated for both imaging days, to quantify the change we took the ratio between both days as 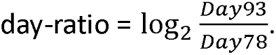

**Table 1:** Vegetation indices used in this study and their formulas. . A more complete version can be found in Supplementary table 2.

| Trait | Full name | Formula |
| --- | --- | --- |
| ARVI | Atmospherically Resistant Vegetation Index | $\frac{NIR - red - (red - blue)}{NIR + red - (red - blue)}$ |
| Clred | Chlorophyll Index Red Edge | $\frac{NIR}{RedEdge}$ |
| EVI | Enhanced Vegetation Index | $2.5 \cdot \frac{NIR - red}{NIR + 6 \cdot red - 7.5 \cdot blue + 1}$ |
| NDRE | Normalized Difference Red Edge Index | $\frac{NIR - RedEdge}{NIR + RedEdge}$ |
| NDVI | Normalized Difference Vegetation Index | $\frac{NIR - Red}{NIR + Red}$ |
| SIPI* | Structure Insensitive Pigment Index | $\frac{NIR - blue}{NIR - red}$ |
| SR | Simple Ratio | $\frac{NIR}{red}$ |
| WDVI | Weighted Difference Vegetation Index | $NIR - a \cdot Red; a = \frac{NIR(soil)}{red(soil)}$ |

\* This index requires light at 800nm, 680nm and 445nm. Since we do not have those exact wavelengths available, we use NIR, red and blue as an approximation.

### Phenotype clustering

Many of the traits we quantify are highly correlated. To make interpreting this number of traits easier we group the traits into 10 k-means clusters based on the square root of the trait*trait correlation matrix with the kmeans() function from base R (R Core Team, 2023). We visualize these clusters as a network using the R package GGnetwork (Briatte, 2023). In this network all traits with an absolute correlation above 0.8 are connected.

### SNP matrix

To genotype all accessions in our study, we combined raw Illumina short read sequencing data of 130 *L. sativa* accessions from Wei *et al*. (2021) with an additional 67 *L. sativa* accessions from van Workum *et al*. (2024a) (European Nucleotide Archive (ENA) at EMBL-EBI, PRJEB63589). SNP calling was done against the Salinas V8 Lettuce reference genome, downloaded from NCBI (GCF_002870075.4, https://www.ncbi.nlm.nih.gov/genome/352)

The BAM files generated in van Workum *et al*. 2024a were used. We added readgroup information to each BAM file to enable using the GATK toolkit (version 4.1.3.0; Poplin et al., 2018). The resulting bam files were converted to .cram format for long-term storage using the following command for each LK sample: samtools view --threads 8 -O cram,embed_ref -T reference.fasta -C -o LK.cram LK.bam The variants were called using the readgroup-added and markduped bam files as input for the variant-calling part of the pipeline: https://github.com/UMCUGenetics/NF-IAP/tree/bqsr_optional_gatk_parameterized First, we ran the GATK HaplotypeCaller on each sample. The resulting GVCF files were split into smaller genomic chunks. All samples were then combined for each chunk (CombineGVCF). Then the combined per chunk GVCFs are Genotyped by GATK GenotypeVCF. After that, variant-selection and filtration were performed (SNPs and INDELs). The results were then again combined for all samples per chunk. This resulted in a combined VCF containing SNPs and INDELs for all samples per chunk. Lastly, all the separate chunks were merged into a single large VCF file (**Box 1**).

#### Box 1 Bash code to generate SNP map.

The bash commands used to combine SNPs and INDELs into a SNP map to be used for GWAS.

HaplotypeCaller --params.haplotypecaller.optional = ‘-ERC GVCF’

VariantFiltration ‘SNP’ -> “--filter-expression ‘QD < 2.0 || FS > 60.0 || MQ < 40.0 || SOR > 3.0 || MQRankSum < -12.5 || ReadPosRankSum < -8.0’ --filter-name ‘snp_filter’ --genotype-filter-expression ‘DP < 2 || DP > 50’ --genotype-filter-name ‘dp_fail’”

’INDEL’ -> “--filter-expression ‘QD < 2.0 || FS > 200.0 || SOR > 10.0 || MQRankSum < -12.5 || ReadPosRankSum < -8.0’ --filter-name ‘indel_filter’”

SelectVariants ‘SNP’ -> ‘--select-type SNP --select-type NO_VARIATION’ ‘INDEL’ -> ‘--select-type INDEL --select-type MIXED’

The SNPS in the obtained VCF were filtered and processed. SNPs were excluded when they had less than 10 of either reference alleles (REF) or alternative alleles (ALT) or more than 10 absent alleles. Next, SNPs were grouped based on linkage to reduce the size of the data as well as the multiple testing burden. SNPs were inspected for possible groups/clumps in batches of 2000 SNPs. Starting at the first SNP in the list of all SNPs and the first of each following batch of 2000 SNPs, each following SNP that differs from the initial SNP in less than 5 positions, out of 198, were grouped. These SNPs, the SNP under inspection and the SNPs grouped to that SNP) were discarded from the list. Then a new first SNP was considered, and so on, the process repeated until all SNPs were assigned to a group. Positions of each SNP within a group were recorded and can be traced. The allele distribution of the SNP with the highest MAF was taken as the representative. The resulting matrix, with 0 for REF, 1 for HET and 2 for ALT, was used for GWAS.

### GWAS

GWAS was performed on all traits using R (version 4.3.1) (R Core Team, 2023) and the Ime4QTL package (Ziyatdinov *et al*., 2018). SNPs were obtained from the raw sequences from Wei *et al*. (2021) and from newly sequenced lines (Van Workum et al., 2024a). The SNP covariance matrix (R-function cov(), **Supplementary Table 4**) was used to correct for population structure. Taking a conservative approach to multiple testing correction, we only considered SNPs with a - log_10_ *pvalue* > 7 significantly linked to a trait. Broad-sense heritability for each trait was calculated by taking the ratio of the between genotype variance and the total variance, using the mean square values as measures of variance.

### QTL selection

To find QTLs, we binned the genome into ∼1Mbp bins and collected the lowest p-value per bin. We defined prominent loci for the mean traits by selecting regions where at least 3 bins in a window of 5 mega-basepairs (Mbp) had a SNP with a - log_10_ (p) score > 8. The lower and upper boundary of the regions was determined by searching for the most distant SNPs with a score of at least 8 in 20 Mb window around the SNP with the highest score. For the extended descriptives we applied stricter criteria. In this case, we selected any locus within a 5 Mbp window, that had at least one SNP with - log_10_ (p) > 12 and at least two other SNPs with - log_10_ (p) >11, and at least three unique traits in the window. The threshold for the boundaries was set at - log_10_ (p) > 11. When looking for candidate genes near the most significant SNPs we use the gene annotation from Van Workum et al., 2024b).

### Code and data availability

Scripts used for this study are available on github: https://github.com/SnoekLab/Dijkhuizen_etal_2025_Drone.

Extended data available on https://doi.org/10.24416/UU01-S5FCM9. This includes all raw data to reproduce results, all intermittent steps, the data required to generate all figures and the weather data.

## Results

### Phenotypic variation

To investigate the genetic basis of color and height traits in lettuce we grew 194 lettuce accessions (**Supplementary Table 1**) near nature area “de Peel” in the Netherlands (**Figure 1**). All data was obtained with the help of an unmanned aerial vehicle (UAV). Drone phenotyping was done on the 11^th^ of June 2021 (78 days after sowing) and on the 25^th^ of June 2021 (93 days after sowing). We quantified 75 traits from this data (**Supplementary Table 5**). These traits consist of the RGB and MSP values, several common vegetation indices, and height. We also used several color ratios as traits (**Supplementary Table 2**). First, we performed GWAS on the mean traits and later we investigated the added benefit of using other descriptives such as the quantiles or standard deviation.

Substantial variation in these 75 traits was observed across the seven different lettuce morphology types, both within and between days (**Figure 1**, **Figure 2, Supplementary Figure 2, 3)**. For example, we noticed that cos, cutting, and stalk lettuce types had the largest increase in height between the two measuring days, whereas the oilseed type had the smallest increase (**Figure 2A**). The red edge trait, which describes the reflection on the edge of visible to infra-red light, was found to increase from day-78 to day-93. Oilseed types had the lowest values whereas butterhead types had the highest values, especially at day-78 (**Figure 2B**). The green/blue ratio, which describes the color of the leaves was found to increase from day-78 to day-93 for almost all accessions. The purple-colored accessions reflect more blue than green light (negative values in **Figure 2C**). Most purple-colored accessions are found in cutting lettuce. In summary, the lettuce population shows substantial variation within and between measurement days and within and between lettuce types.

**Figure 2:**
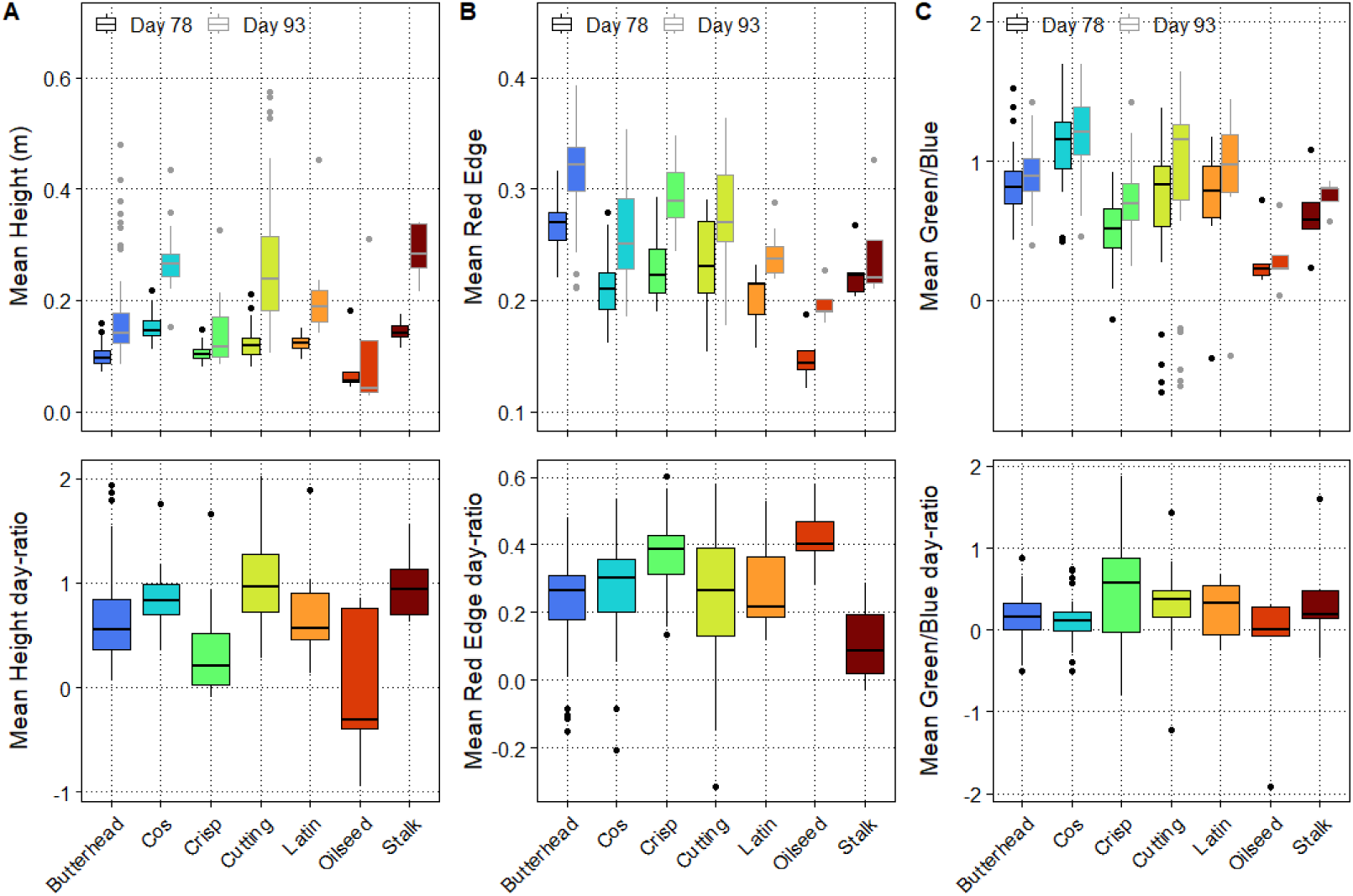
Variation in selected traits between seven lettuce types over two timepoints. **A**) Mean height of seven lettuce types over two time points **B**) Mean Red Edge values from the Multispectral camera **C**) Mean Green/Blue ratio from the RGB camera. Upper panels show the values at day-78 and day-93 and lower panels show the day-ratio. Lettuce types are indicated on the x-axis and by the different colors.

Most of the phenotypic variation was found to be highly heritable (**Supplementary Figure 4, 5, Supplementary Table 6**). The average heritability on day-78 and day-93 is 82.7% and 84.1% respectively. Whereas the day-ratio heritability is slightly less to hardly heritable with an average heritability of 61.4%. The minimum and skewedness had the lowest heritabilities out of the different trait descriptions. In contrast, the maximum and high quantiles often show increased heritability compared to the mean trait values. Overall, these 194 lettuce varieties displayed considerable heritable phenotypic variation, enabling investigation of the genetic architecture of these traits.

### Correlation between traits

Traits that are highly correlated are redundant and would likely map to the same QTLs. A high correlation, both positive and negative, likely indicates a shared genetic architecture. To quantify the similarity between the traits, we calculated the Pearson correlation between all traits (**Supplementary Table 11**). Many traits showed high correlation, such as height at day-78 and NDVI at day-78. To visualize this, we constructed a correlation network with traits as nodes and correlations exceeding 0.8 as edges (**Figure 3**). Most (68 of 75) of the traits share edges and can be found in clusters. The rest of the traits (7 of 75) are isolated from the rest of the network. Some just fall short of the correlation threshold. Others, such as the change in Red-blue ratio are not correlated to any of the other traits (**Supplementary Table 11**). We used K-means, an unsupervised clustering method, to divide the network into 10 clusters based on the squared correlation matrix of the traits. (**Figure 3)**. We then named all these clusters based on the most abundant traits in the cluster to make discussion and interpretation easier (**Table 2**). In general, traits from day-78, day-93, and especially the traits describing the day-ratio tend to correlate more within their respective groups than between groups. We also saw that the relative blue (**Figure 3, cluster F**), Red- /green (**cluster E**) and Relative red cluster (**cluster I**) are connected in the network. This was as expected as all these traits are a description of how red, purple, or green the leaves are. Other large clusters were cluster B, formed entirely by color ratio traits, cluster C, consisting mostly of several vegetation indices aimed at quantifying chlorophyll content and cluster D, formed by several traits relating to the greenness of the plant. Clusters of traits likely indicate that these traits share part of their genetic architecture and are partial descriptions of a more complex phenotype. Due to the clusters of correlating traits we expected to find several shared loci using GWAS for traits in the same cluster, as opposed to a unique SNP association pattern for each individual trait.

**Figure 3:**
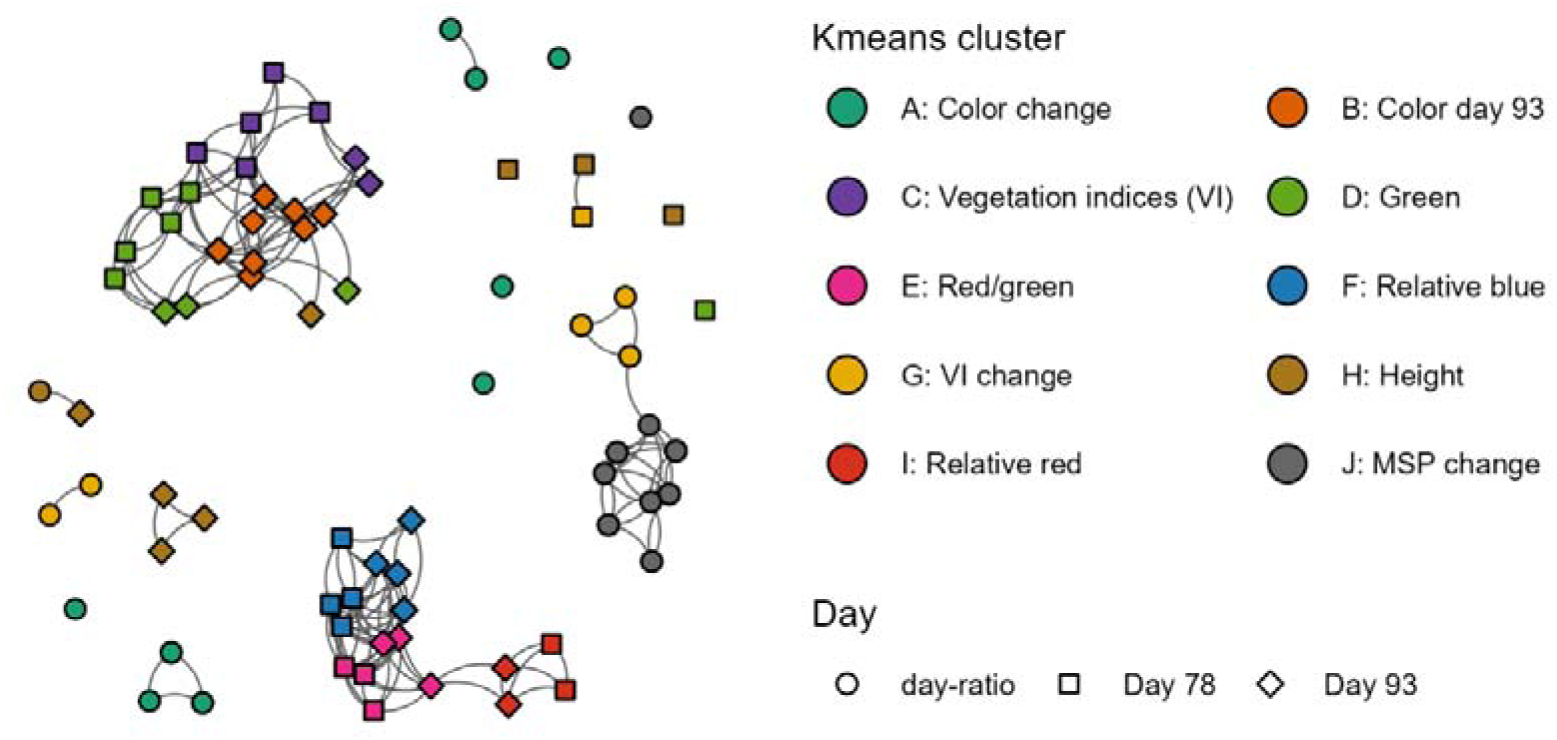
Visualization of correlation between traits using clusters obtained using K-means clustering. K-means clustering on all traits. Each node represents a trait, and each edge represents a correlation > 0.8 between traits. Trait observation type, day-78, day-93 and day-ratio are shown by the different shapes. The K-means clusters are indicated by the different colors shown in the legend.

**Table 2:**
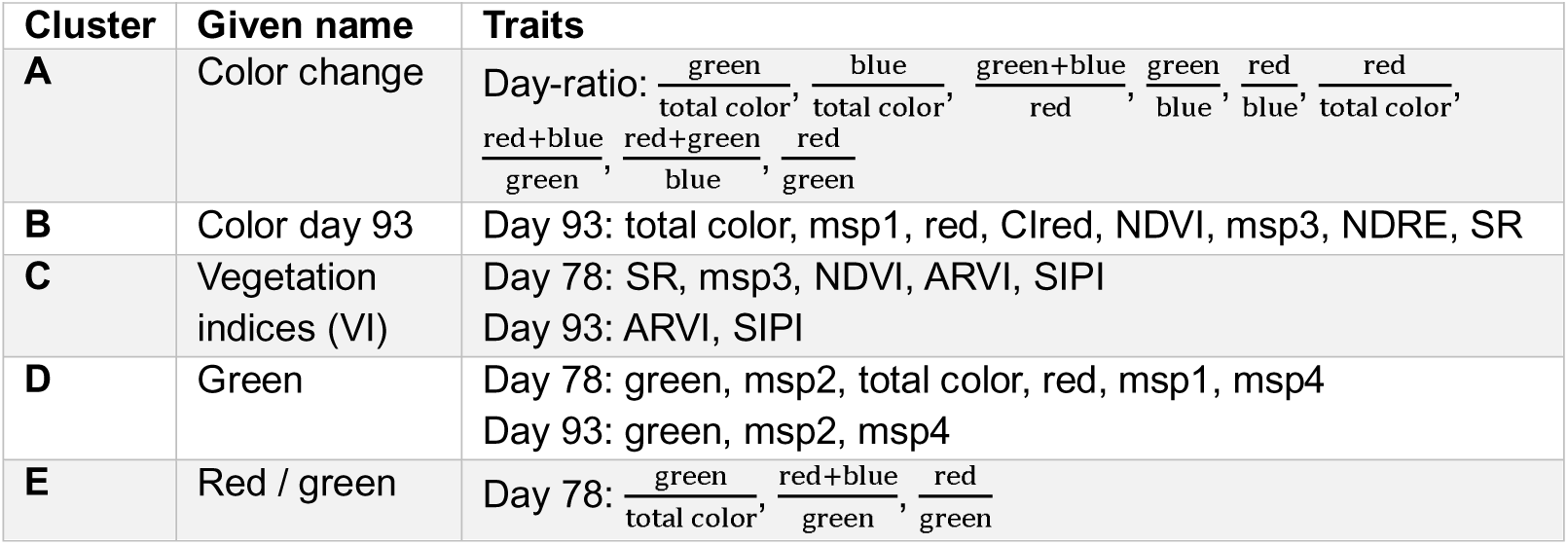
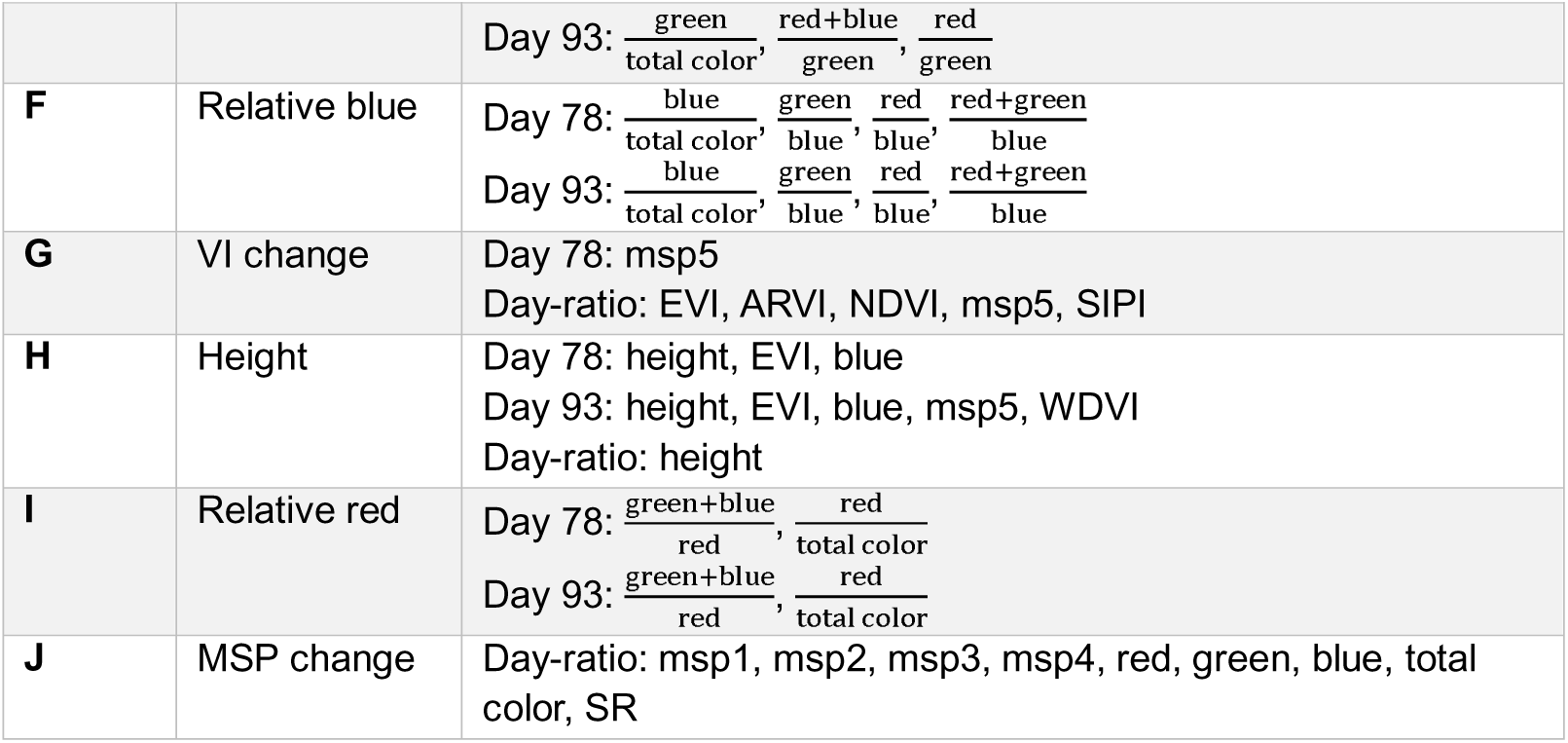
Contents of all trait clusters. All traits for clusters A:J sorted by which day the traits were measured. All clusters were given a name that best represents their contents to make discussion easier.

### Genome wide association

To uncover the polymorphic loci underlying the observed phenotypic variation, we performed SNP-based GWAS for all 75 traits. We found significantly associated SNPs (- log_10_ (p) > 7) for 55 traits (**Table3**). As examples we show the GWAS results for the day-93 Green, Red Edge, and Height traits. (**Figure 4, Supplementary Table 7,8,9**). For the variation in height at day-93 we found a QTL on chromosome 7 at 164.4 megabasepairs (Mbp). The QTL locus contained the previously identified phytochrome C locus associated with bolting and flowering time in lettuce (Rosental *et al*., 2021). This QTL is not detected when using the height at day-78 as a trait for GWAS (**Supplementary Figure 6**). This result suggests that a substantial number of lettuce varieties started bolting between the first and second day of phenotyping. The green to blue ratio describes whether the color is more green or more purple. This is a good proxy for anthocyanin content. For this trait we recover *RLL2* (chr 5, 86.7 Mbp), *RLL4* (chr 9, 95.1 Mbp) and ANS (chr 9, 152.6 Mbp). All these loci are essential to the typical purple color seen in some lettuce varieties (Su *et al*., 2020). When looking at the red edge intensity we find a QTL at the “pale leaf locus” (chr 4, 106.0 Mbp). This locus harbors the *LsGLK* gene, which is involved in variation in chlorophyl content (Zhang *et al*., 2022). Recovering these known loci solidifies the validity of using drones for phenotypic observation.

**Figure 4:**
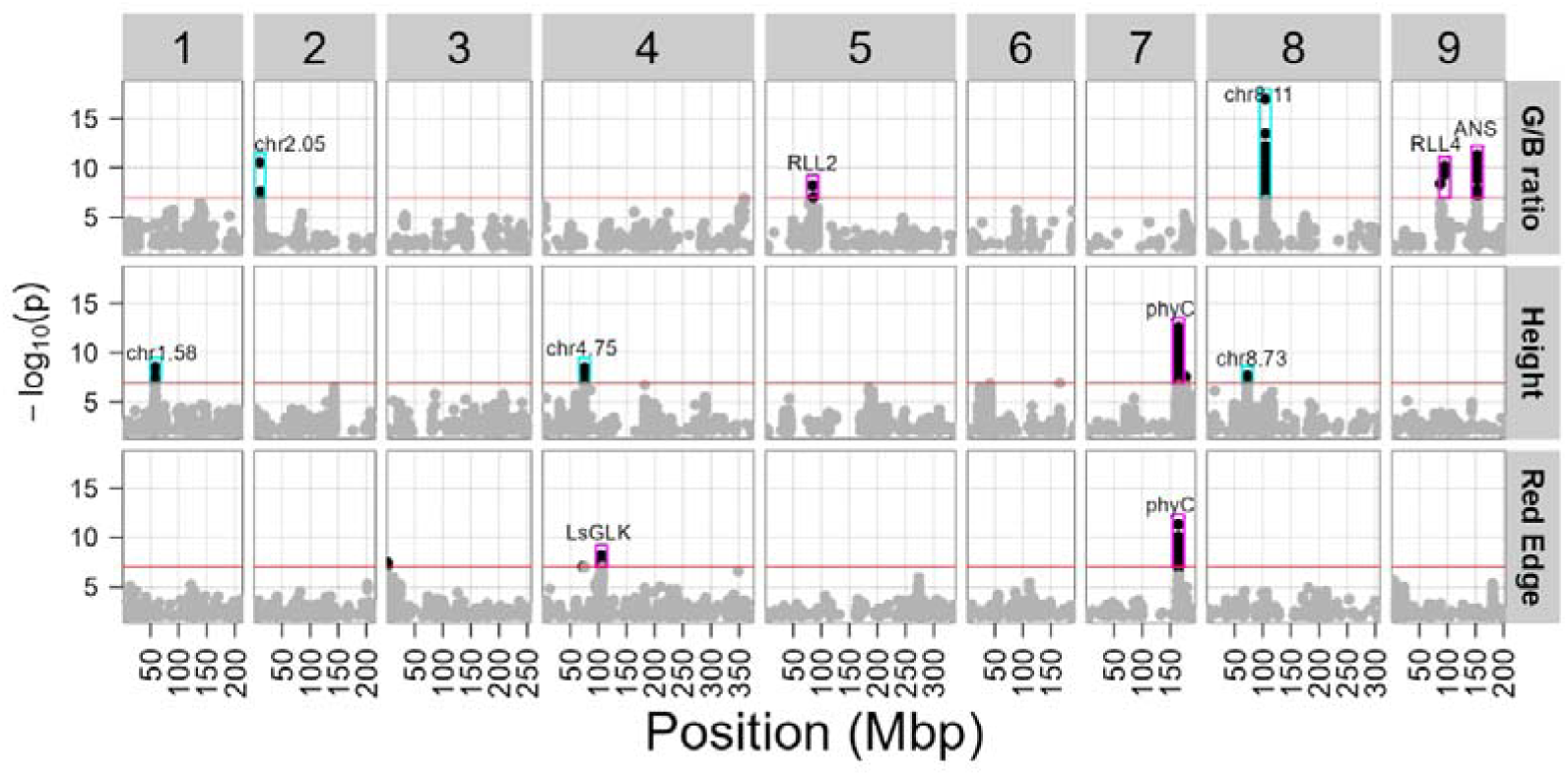
Manhattan plots for G/B ratio, height and Red Edge on day 93 show multiple QTLs. The x-axis represents the genome of *L. sativa* in mega-basepairs (Mbp). Chromosomes are shown on top. The y-axis represents the p-value of the association between SNPs and the specific trait. The red line at -log_10_(p-value) = 7 represents the multiple testing significance threshold. SNPs above this threshold are shown in black and those below are shown in grey.

To visualize the combined GWAS results of all 55 traits with significant SNPs, we aggregated all SNPs (-log10(p) > 7) into one figure (**Figure 5, Supplementary Table 10**), showing each cluster separately. High confidence QTLs were selected by taking all loci where at least three SNPs have a -log_10_ (p) score > 8 within 5 Mbp of each other (**Table 3, Supplementary Table 12, cyan and purple rectangles in Figure 5**). We hypothesized that traits that clustered together based on correlations would have a shared genetic basis. Our results substantiate this hypothesis, since every cluster has its own pattern of detected QTLs. Furthermore, clusters that grouped together in the correlation network **(Figure 3**) also had overlapping QTLs. For example, the Relative blue (F), Red/Green (E) and Relative red cluster (I) grouped together phenotypically and showed similar GWAS results. The Green cluster (D) was an exception, correlating mostly with the Color day-93 (B) and VI (C) clusters but showing similar GWAS results to the Red/Green (E) and Relative blue (F) cluster. Many QTLs were recovered by several traits from different clusters, such as the QTLs containing *RLL2*, *RLL4*, Anthocyanidin synthase and *phyC*. This aligns with expectations as many of the color traits show overlap and have a high correlation.

**Figure 5.**
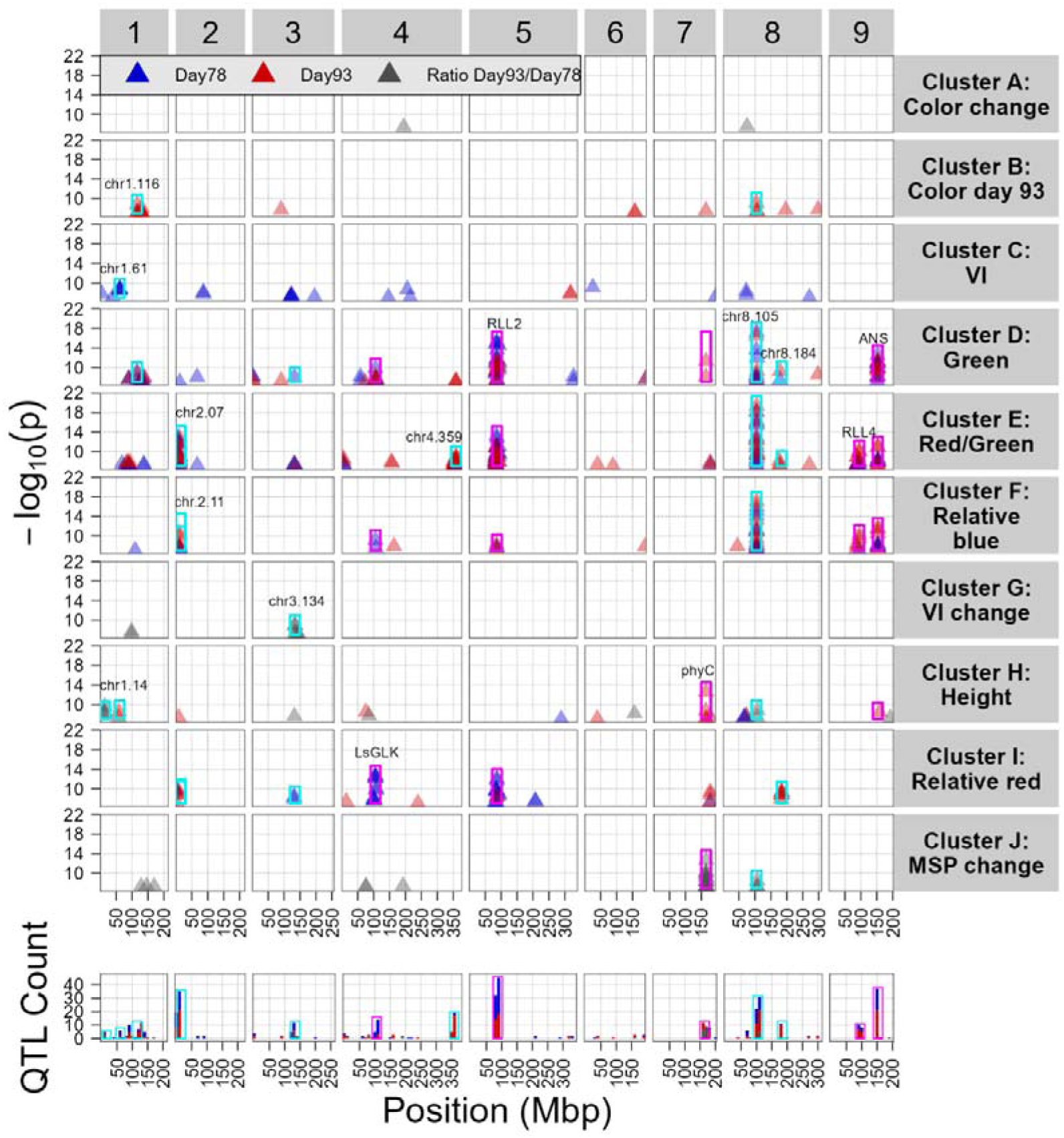
Multi-Manhattan plot of all trait clusters. Every row represents one of the clusters, shown at the right. The y-axis represents the -log_10_ (p-value) of the peak SNPs per trait. Loci that have been previously reported in literature (**Table 3**) are annotated with a purple rectangle and new loci with a cyan rectangle. The color of the points represents whether the traits are from day-78 (blue), day-93 (red), or the difference between the two days (grey). The x-axis represents the position on the genome of *L. sativa* in megabasepairs (Mbp). At the bottom a histogram depicts the total amount of significant (-log_10_(p) > 7) in a 10 Mbp window.

**Table 3:** All prominent QTLs using mean color and height values. From all the GWAS results we selected prominent QTLs (all QTLs can be found in **Supplementary Table 12**). To qualify a locus needs at least three SNPs with a score above 8 within 5 Mbp. We reported the statistics of the trait with the most significant QTL. The first and second columns show the chromosome and position of the most significant SNP of the QTL peak in megabasepairs (Mbp) on the lettuce reference genome v8 (ncbi). The third column shows the SNP’s most significant trait. The final column shows the source from previous literature.

| Chromosome | Position Mbp (v8) | Most significant trait | $-\log_{10}(p)$ max | Candidate gene* | Source candidate gene |
| --- | --- | --- | --- | --- | --- |
| 1 | 14.06 | $\frac{\text{Red}}{\text{TotalColor}}$ day-ratio | 9.59 | | |
| 1 | 60.95 | SIPI day-78 | 8.82 |  |  |
| 1 | 115.83 | Green day-78 | 9.03 |  |  |
| 2 | 0.70 | $\frac{\text{Red}}{\text{Green}}$ day-78 | 13.25 | | |
| 2 | 11.24 | $\frac{\text{Red} + \text{Blue}}{\text{Green}}$ day-93 | 12.92 | | |
| 3 | 133.85 | NIR day-ratio | 8.97 | *GST* | (Zhang *et al.*, 2017) |
| 4 | 106.00 | $\frac{\text{Green} + \text{Blue}}{\text{Red}}$ day-78 | 12.67 | *LsGLK* | (Zhang *et al.*, 2022) |
| 4 | 358.95 | $\frac{\text{Red}}{\text{Green}}$ day-93 | 9.09 | | |
| 5 | 86.67 | Green day-78 | 15.34 | *MYB113/114 (RLL2)* | (Su *et al.*, 2020; Zhang *et al.*, 2023) |
| 7 | 165.02 | Red Edge day-ratio | 12.69 | *phyC* | (Rosental *et al.*, 2021) |
| 8 | 105.48 | $\frac{\text{Red} + \text{Blue}}{\text{Green}}$ day-93 | 19.33 | | |
| 8 | 183.81 | $\frac{\text{Red}}{\text{TotalColor}}$ day-93 | 9.44 | | |
| 9 | 95.11 | $\frac{\text{Red} + \text{Blue}}{\text{Green}}$ day-93 | 10.22 | *RUP1/2 (RLL4)* | (Su *et al.*, 2020) |
| 9 | 152.57 | Green day-78 | 12.50 | *ANS* | (Zhang *et al.*, 2017) |

\* For all candidate genes at the locus see Supplementary table 12. This table includes several suggestions for the novel QTLs.

On top of the previously known QTLs, we also found several new QTLs that to our knowledge have not been reported before in lettuce. For height at day 93 we found QTLs on chromosome 1 at 58.5 Mbp, chromosome 4 at 75.4 Mbp, and chromosome 8 at 72.8 Mbp (**Figure 4**). Strikingly, we found a novel QTL with high confidence (-log10(P) > 19) on chromosome 8 at 105 Mbp for the green to blue ratio and several other traits relating to anthocyanin content. This QTL has not been reported in lettuce for any anthocyanin related trait before. Most accessions that are homozygous for the alternate allele at the closest locus show the typical purple color associated with anthocyanin (**Supplementary Figure 7**). We noticed the candidate gene *FQR1* (105.00 Mbp) close to the most significant SNP (105.48 Mbp). This gene is described as a flavodoxin. Although there is no direct relation between flavodoxins and flavonoids this FQR1 protein has already been shown to affect pigmentation in vitro in *Arabidopsis* (Laskowski *et al*., 2002) making it a feasible candidate gene. We found another QTL on the same chromosome (chr8, 183,81 Mbp) for several traits relating to the green-red ratio **(Supplementary table 10)**.

### Extended descriptives

In addition to analyzing the mean of the traits, we also investigated several extended descriptives to describe different aspects of certain phenotypes, potentially yielding distinct GWAS results. The extended descriptives we used are trimmed means, several quantiles, the median, the standard deviation, the minimum and maximum value, and the skewness and kurtosis of the trait distribution (**Supplementary Table 3**). To test the usefulness of these descriptives, we compared them to the GWAS results obtained using only the mean trait values. We hypothesized that these descriptives are more sensitive to specific phenotypes than the mean trait value. For example, when a lettuce plant starts bolting this should have a much stronger effect on the maximum height of the plant than the average of plant parts together (**Figure 6A**,)**B**. To test this, we compared the results of all mean height traits with their extended descriptives (**Figure 6C, Supplementary Table 14**). We examined both the day-ratio and the absolute difference between days.

**Figure 6.**
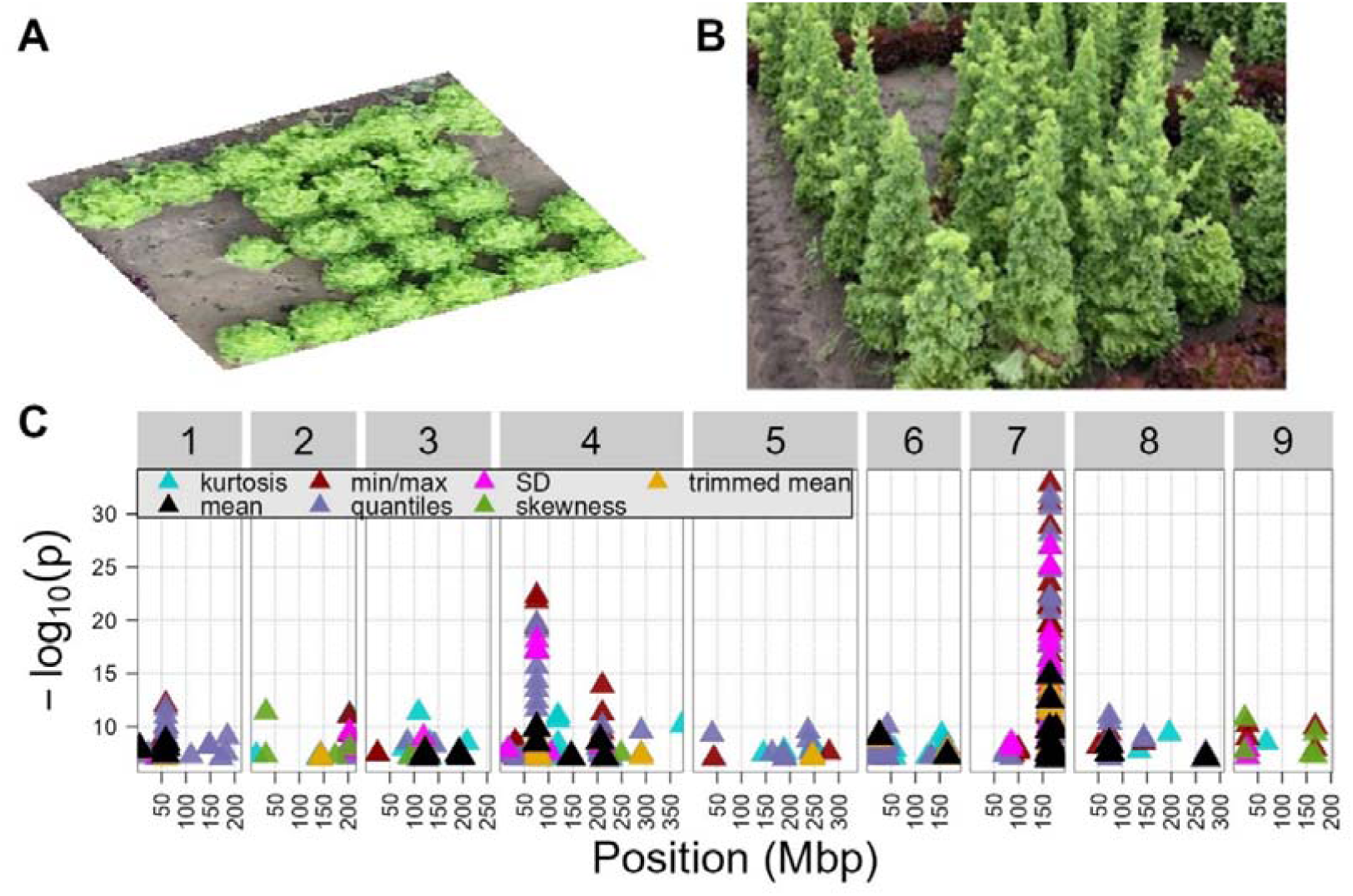
GWAS on traits related to the change of height between day-78 and day-93. A) A lettuce variety bolting as seen from a top perspective. B) A bolting lettuce variety as seen from a side perspective. C) The GWAS results on all height traits. In black the results of the GWAS on the mean traits and in colors several extended descriptives.

We found that the extended descriptives sometimes represented the traits better than the mean, leading to higher significance in the GWAS. For example, the mean height traits mapped to the *phyC* locus with a -log_10_ (p-value) of 12.6, yet the maximum height ratio between day-93 and day-78 maps to the same locus with a -log_10_ (p) score of 32.9 (**Figure 6C)**. Similarly, both analysesmapped to a QTL on chromosome 1 60.0 Mbp with a -log_10_ (p) score of 8.5 for mean height, whereas the maximum and quantiles of height mapped to this QTL with a higher -log_10_ (p-value) of 12.1 (**Figure 6C, Supplementary Tables 12,13**). Additionally, we uncovered loci not identified when using the mean traits. For example, we recovered a highly significant (-log_10_ (p) = 22.3) QTL for height at chromosome 4 at 75.5 Mbp that fell short of our strict definition for a prominent QTL when only using mean traits (**Figure 6C**).

Following the same strategy of clustering traits we performed a new k-means clustering, combining the mean traits with all extended descriptives (**Supplementary Figure 8**). The clustering on all traits was remarkably similar to the clustering on only mean traits. Here we also found a height cluster, a red cluster, and a green cluster (**Supplementary Figure 9**). Most QTLs found with the extended descriptives were also found with the mean traits (**Supplementary Figure 8**). Notably, using extended descriptives as traits in GWAS resulted often in detecting these shared QTLs with a higher significance. Using extended descriptives (the maximum green reflection), we found the previously reported *RLL3* locus (Su *et al*., 2020), which we did not find using only mean traits (**Supplementary Figure 8, Supplementary Table 13**). This shows that the utilization of quantiles and maxima is beneficial for identifying novel QTLs. However, the standard deviation, skewness and kurtosis appear to give unreliable results as these descriptives have a low heritability and fail to uncover clear loci (**Supplementary Figure 5, 10:19**). In summary, extended descriptives are often better trait descriptions compared to the mean leading to higher confidence mapping and additional QTLs.

## Discussion

In recent years, advancements in genotyping technologies have alleviated the bottleneck previously posed in GWAS studies (Spindel *et al*., 2018; Watanabe *et al*., 2017). Now, phenotyping is often considered to be the limiting step, especially in field conditions. In this study we grew 194 lettuce accessions in a field setting and employed drone-assisted imaging and automatic phenotyping to unravel the genetic underpinnings of color and height traits. Our GWAS uncovered several previously known QTLs and several new QTLs associated with these traits. By using descriptives other than the mean such as quantiles, minima and maxima we detected many loci with a higher confidence and uncover new loci. In addition, we showed that by using remote sensing to non-destructively phenotype multiple days we can uncover genetic loci that have a stronger effect on trait variation in one developmental stage compared to another.

### Previously discovered QTLs

Multiple previous studies have investigated the genetic background of color and height traits in lettuce (Wei *et al*., 2021; Su *et al*., 2020; Falcioni *et al*., 2023; Rosental *et al*., 2021). Thus, some of the QTLs we found have been discovered before. Our analysis identified QTLs at *RLL2*, *RLL3*, and *RLL4* (Zhang *et al*., 2023; Su *et al*., 2020) and ANS, the latter being responsible for the pen ultimate step during anthocyanin synthesis (Wilmouth et al., 2002; L. Zhang et al., 2017). These QTLs are associated with several traits describing the purple color in the plant, which is a proxy for anthocyanin content. We also uncover a QTL containing *LsGLK*, a gene that plays a pivotal role in chloroplast development. This gene was previously shown to be vital for the typical green color present in many lettuce varieties (Chen et al., 2016; L. Zhang et al., 2022). We also identify a QTL containing the well-known bolting gene *Phytochrome C* (Rosental *et al*., 2021), for color and height traits, especially height on day-93 and day-ratio. A previous study investigating flowering time and anthocyanin content in lettuce under field conditions mapped QTLs containing *phyC*, *RLL2* and *ANS* (Wei *et al*., 2021). We reproduce these QTLs as well as QTLs containing *RLL3, RLL4,* and several novel QTLs. There are a few possible explanations for the fact that our study finds additional QTLs compared to the study by Wei *et al*. One explanation is simply a difference in population, causing additional segregating alleles affecting color and height traits. Another possible reason is that many QTLs are only found at specific developmental stages or in certain environmental conditions (Bac-Molenaar *et al*., 2016; van Eijnatten *et al*., 2024; Vasseur *et al*., 2014). The study from Wei *et al*, which measured only one timepoint, may have missed some QTLs whose effects depend on the developmental stage. This might also explain why we do not find a QTL containing *RLL1*, a gene that was previously shown to affect anthocyanin content and color traits (Su *et al*., 2020). A final explanation of the observation that we discover more QTLs than Wei *et* al is the use of extended descriptives, which helps capture different facets of phenotypic variation. We would not have been able to detect the QTL containing *RLL3* in this experiment using only the mean traits.

### Novel QTLs

In addition to uncovering several previously reported QTLs our analysis uncovered novel QTLs. For several novel QTLs we identified candidate genes by examining the *Arabidopsis* homologs near the QTLs (**Supplementary Table 12**). On chromosome 4 at 75.5 Mbp we find a novel QTL in a GWAS population related to height and height day-ratio. Many of these traits also map to phytochrome C (**Figure 4, Supplementary Figure 6, Supplementary table 13**). Close to this QTL are 2 potential candidate genes. The closest candidate gene to the top SNP (76.2 Mbp) is a homolog of VOZ1 a gene known to promote the flowering transition in *Arabidopsis* (Celesnik *et al*., 2013). Another potential candidate located at 77.1 Mbp is a homolog of *SPL3*, another gene known to affect flowering transition in *Arabidopsis* (Yamaguchi *et al*., 2009). This QTL on chromosome 4 related to height traits potentially overlaps with a previous QTL identified for flowering time in a RIL population (Han, Truco, *et al*., 2021).

We identified a QTL on chromosome 8 at 105.48 Mbp with high confidence (**Figure 5**). This QTL is mostly linked to traits relating to the blue to green ratio on day 93. Examining the allele distribution of the most significant SNP in this QTL, we noticed that most genotypes that are homozygous for the alternative allele show the typical purple anthocyanin color. This is a clear indication that this allele is involved in anthocyanin distribution, since only a handful of all 194 genotypes have this color. (**Supplementary Figure 7**). A potential candidate gene for this QTL might be the homolog of the *Arabidopsis* gene *FQR1* at 105 Mbp. *FQR1* encodes a flavodoxin which has been shown to affect pigmentation in vitro (Laskowski *et al*., 2002). Another candidate gene in this QTL is a homolog of the R2R3 MYB transcription factor MYB119. This transcription factor is known to be involved in the regulation of anthocyanin production in *Arabidopsis*(Cho *et al*., 2016)(**Supplementary Table 12**).

### Timepoints

We conducted imaging at 78 and 93 days old. We imaged twice because we expected different genes to be relevant depending on the developmental stage of the plant. We found that, as expected, some QTLs were unique to traits from either day-78 or day-93. For example, the QTL containing the *LsGLK* gene is almost exclusive to traits from day-78. Conversely, the QTL containing the *phyC* gene is completely exclusive to traits from day-93 and the ratio between the two days. We expected to find the *phyC* gene when looking at the height day-ratio. However, the *phyC* locus is also a QTL for traits like the red-edge reflection and suggests that bolting influences plant-color. Furthermore, we discovered a QTL exclusive to a few day-ratio traits on chromosome 1 at 14.1 Mbp, and a QTL exclusive to traits from day-93 on chromosome 4 at 359.0 Mbp. These results show that traits are impacted by different loci based on the growth stage of the plant. This observation is especially striking since the two imaging days are only 15 days apart, which is a relatively short time span for lettuce growth and development. With the rise of automatic phenotyping, this kind of multi-day-, multi-phenotype-data is likely to become more abundant. This will enable the discovery of more developmental stage-dependent-QTLs in the near future.

### Environment

Apart from variation in development, variation in response to environment can also cause QTL variation. The plants were transplanted outside on the 27th and 28th of April and imaged on the 11th and 25th of June. The plants experienced substantial variation in environment during this period. For example, there was a considerable amount of rain in the two weeks before the second measurement day, noticeably changing the moisture saturation of the soil. The temperature also fluctuated over the course of these two weeks (weather data available on https://doi.org/10.24416/UU01-S5FCM9). Potentially, adaptations to precipitation or temperature could affect the color or height of the plants. Variation between the genotypes in these adaptations could result in QTL that are only found when the environment is in a certain state. Hence, QTLs that switch on or off over time can result from the developmental program, the response to the environment, or the interaction between these.

### Extended Descriptives

In this study we expanded the conventional trait analysis by incorporating various descriptives beyond the mean: the minimum and maximum values, standard deviation, skewness, kurtosis and several quantiles. We have not found any other studies describing traits in this manner. Some of the reported loci would not have been uncovered without these extended descriptives. Especially the maximum and many of the low or high quantiles proved useful to recover known QTLs with higher confidence or even uncover completely new QTLs. This could be explained by these descriptives being more sensitive to spatial variation in leaf color or plant height than the mean or median. The skewness, minimum, and kurtosis showed lower heritability than the other descriptives (**Supplementary Figure 5**). Although these descriptives did result in some significant SNPs, these usually did not form clear peaks of linked SNPs high confidence, (**Supplementary Figure 6:17**). The skewness, minimum or kurtosis were therefore less useful for GWAS analysis. Together, our results show that using extended descriptives can help to discover more relevant genetic variation.

## Conclusion

In summary, we used UAV phenotyping to perform GWAS on plant-color and height in lettuce in a large field experiment. We confirmed several previously known QTLs in field conditions and identified several novel QTLs. We showed that phenotyping at multiple timepoints can uncover development-dependent QTLs. Additionally, we demonstrate the advantage of using extended descriptives such as quantiles and maximum to describe phenotypes, revealing several QTLs that would be missed if we relied only on mean trait values. Our findings highlight that non-destructive phenotyping using a drone can reveal the interactive effect of genes, environment and development on plant color and height traits under field conditions.

## Supporting information

Excel file with necessary data to generate all figures.

## Author contributions

Remko Offringa, Marcel Proveniers, Guido van den Ackerveken and Basten L. Snoek conceived the study. Kiki Spaninks, Remko Offringa, Marcel Proveniers, Guido van den Ackerveken, Sarah L. Mehrem, Jelmer van Lieshout, Esther van den Bergh, and Steven Kaandorp performed the experiment. Abraham L. van Eijnatten, Rens Dijkhuizen, Remko Offringa, Sarah L. Mehrem, and Basten L. Snoek extracted the data and performed the investigation. Rens Dijkhuizen, Abrahm L. van Eijnatten, and Basten L. Snoek wrote the manuscript with input from all co-authors.

## Acknowledgements

We thank the Netherlands Plant Eco-phenotyping Centre (NPEC) for their help with operating the Drone and the first steps in data exchange. We also thank Flip Mulder from the Utrecht Bioinformatics Expertise Core (UBEC) for help with generating the VCF.

## Supplementary material

## Supplementary tables

**Supplementary table 1: LK_lines**

**Supplementary table 2: Pheno_definitions**

**Supplementary table 3: Statistical_definitions**

**Supplementary table 4: kinship**

**Supplementary table 5: Phenotypes**

**Supplementary table 6: Heritability**

**Supplementary table 7: Height.mean.day2**

**Supplementary table 8: gbrat.mean.day2**

**Supplementary table 9: rededge.mean.day2**

**Supplementary table 10: pvalues**

**Supplementary table 11: Correlation matrix**

**Supplementary table 12: QTLs.fig5**

**Supplementary table 13: QTLs.all**

**Supplementary table 14: height.traits**

## Supplementary figures

**Supplementary Figure 1.**
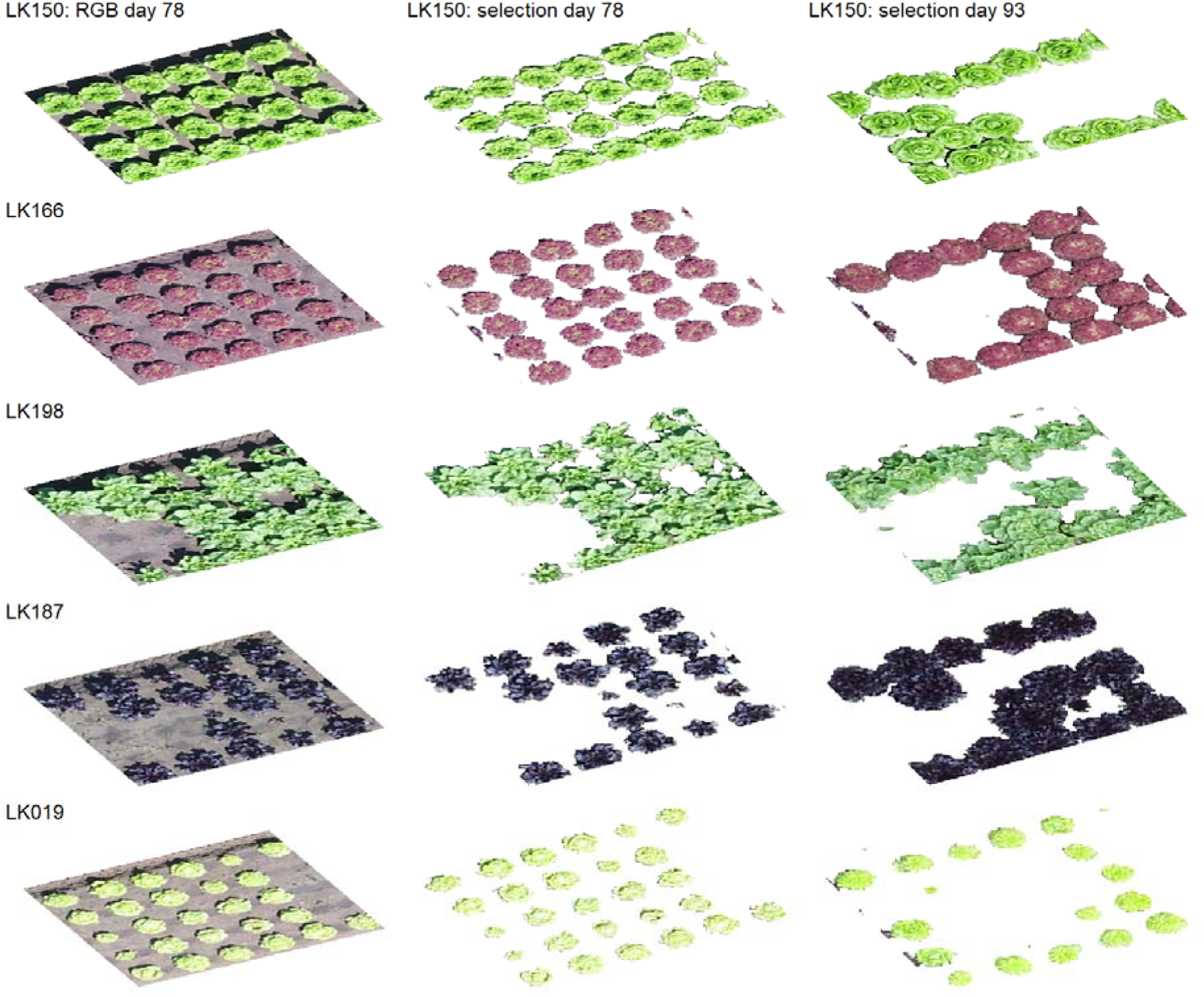
Plant pixel selection using EVI threshold for five accessions. The first column shows the RGB image for the accessions on day 78. The second column shows the plant pixels after thresholding on day 78 with an EVI threshold of 0.25. The third column shows a similar thresholding for day 93 with a threshold of 0.4.

**Supplementary Figure 2:**
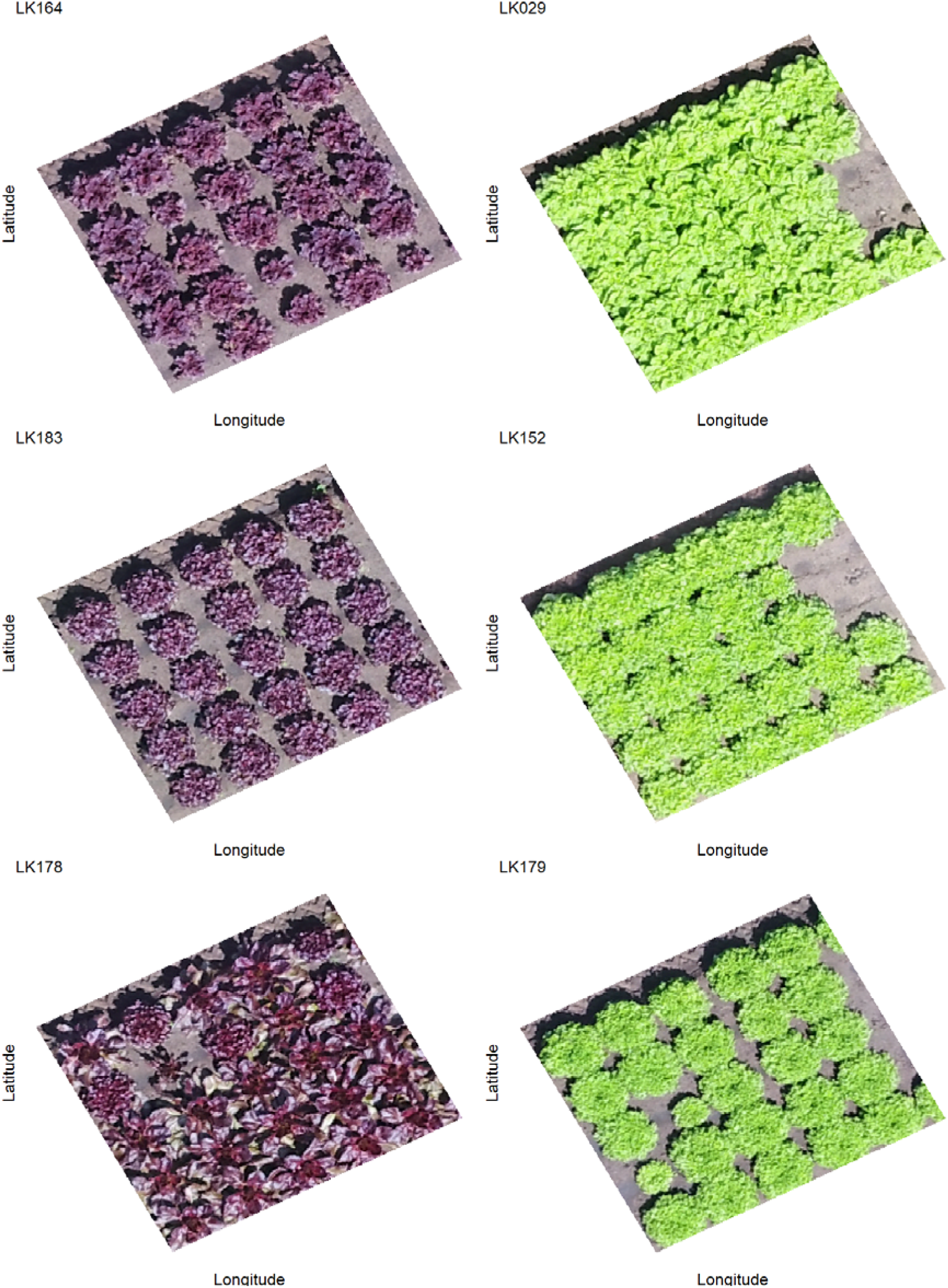
Three accessions with extremely low green/blue ratio on the left and three accessions with extremely high green/blue ratio on the right.

**Supplementary Figure 3:**
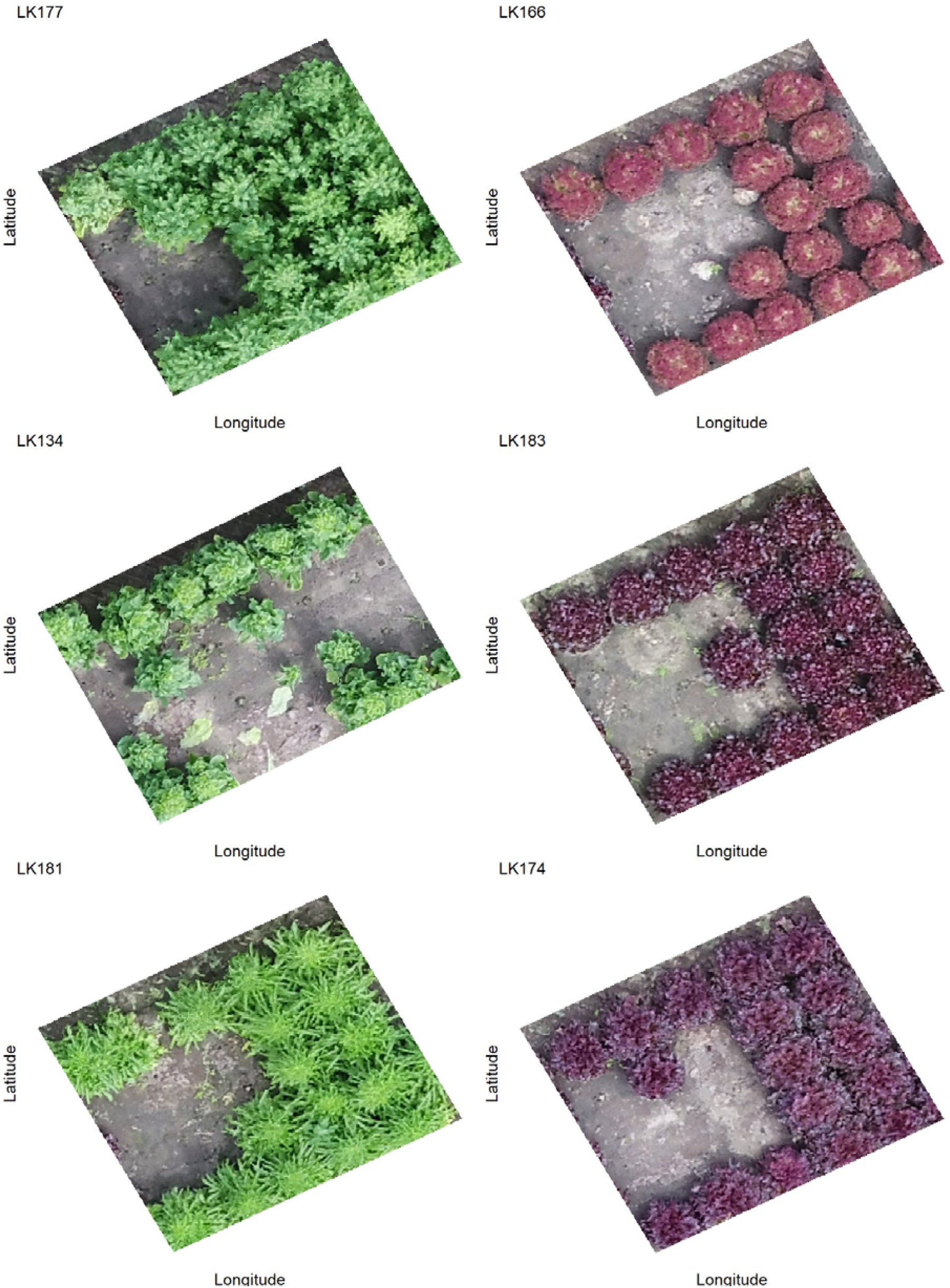
Three accessions with extremely low relative red on the left and three accessions with extremely high relative red on the right.

**Supplement Figure 4:**
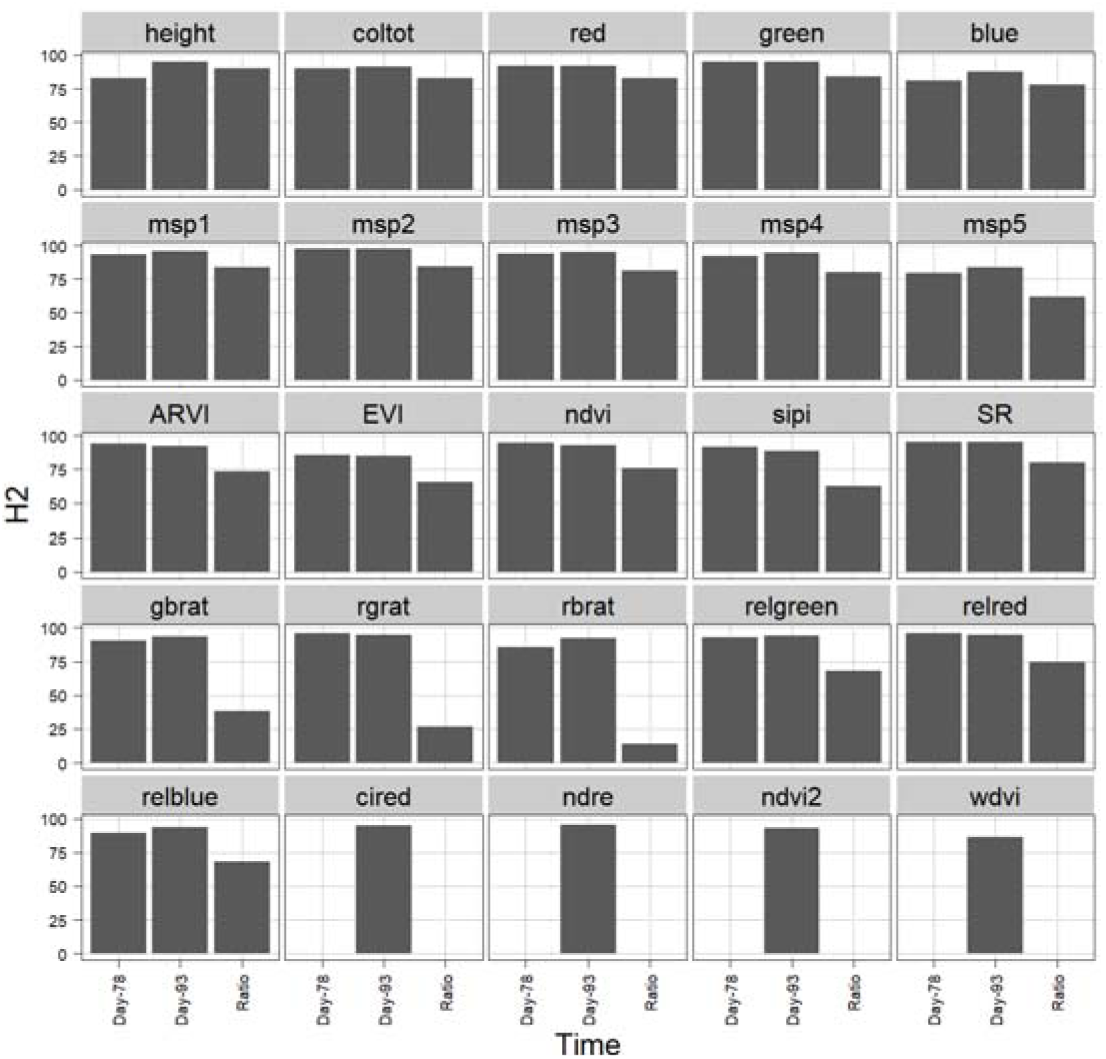
Broad sense heritability of all mean phenotypes. Heritability is shown for day-78, day-93 and the ratio of the trait between those two days.

**Supplement Figure 5:**
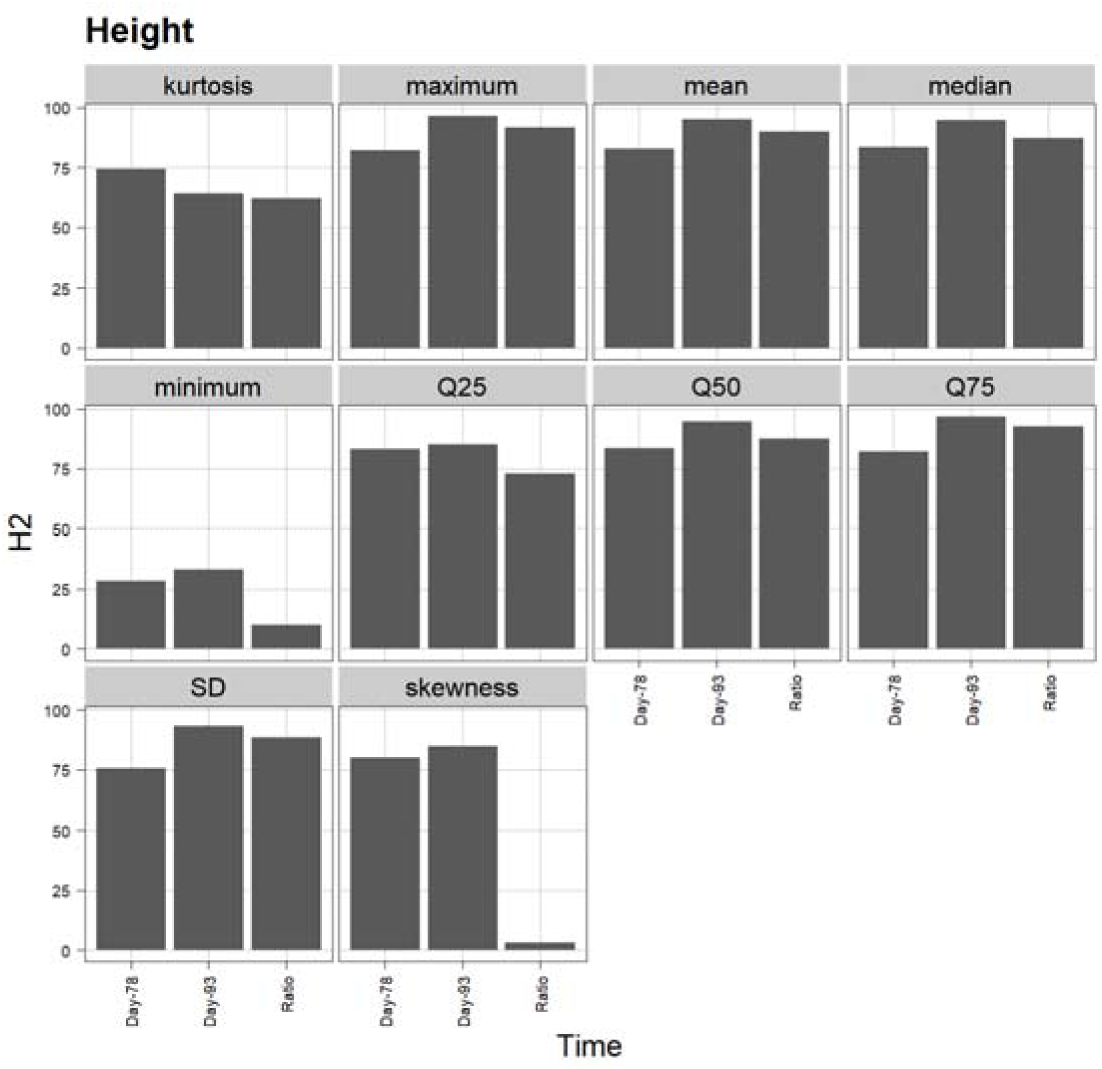
Broad sense heritability of all descriptives of the height trait. Heritability is shown for day-78, day-93 and the ratio of the trait between those two days.

**Supplementary Figure 6:**
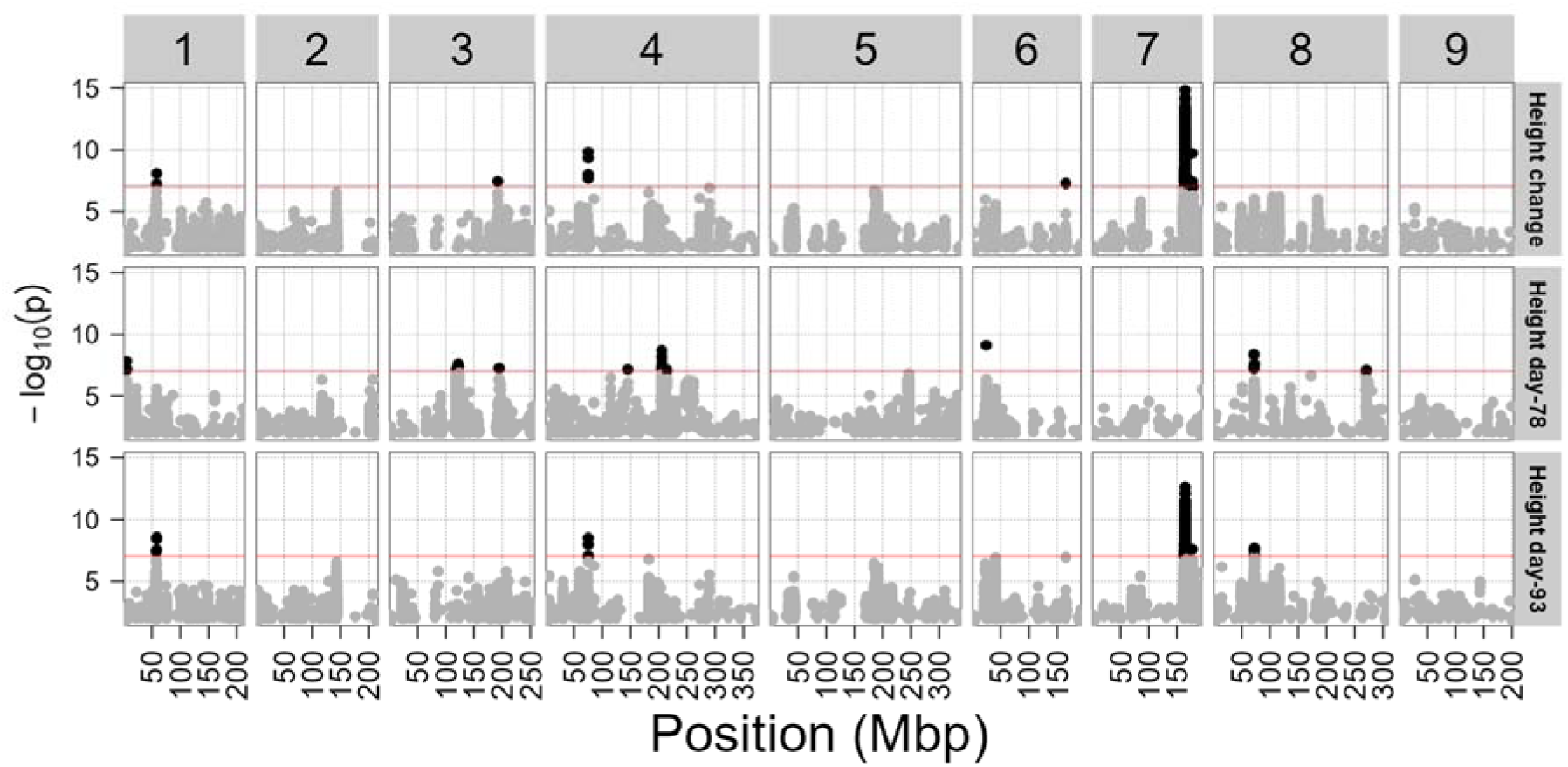
Comparison of height on both days and the day-ratio. The x-axis represents the genome of L. sativa in megabasepairs (Mbp). Chromosome numbers are shown on top. The y-axis represents the p-value of the test associating a SNP is associated to trait variation. The red line at - log_10_(p-value) = 7 represents a conservative significance threshold. SNPs above this threshold are shown in black and SNPs below the threshold are shown in grey. The traits shown here are the height on day-78, the height on day-93 and the absolute change in height between those days. The QTL on chromosome 7 containing Phytochrome C is present on day-93 and the change between both days, but absent on day-78.

**Supplementary Figure 7:**
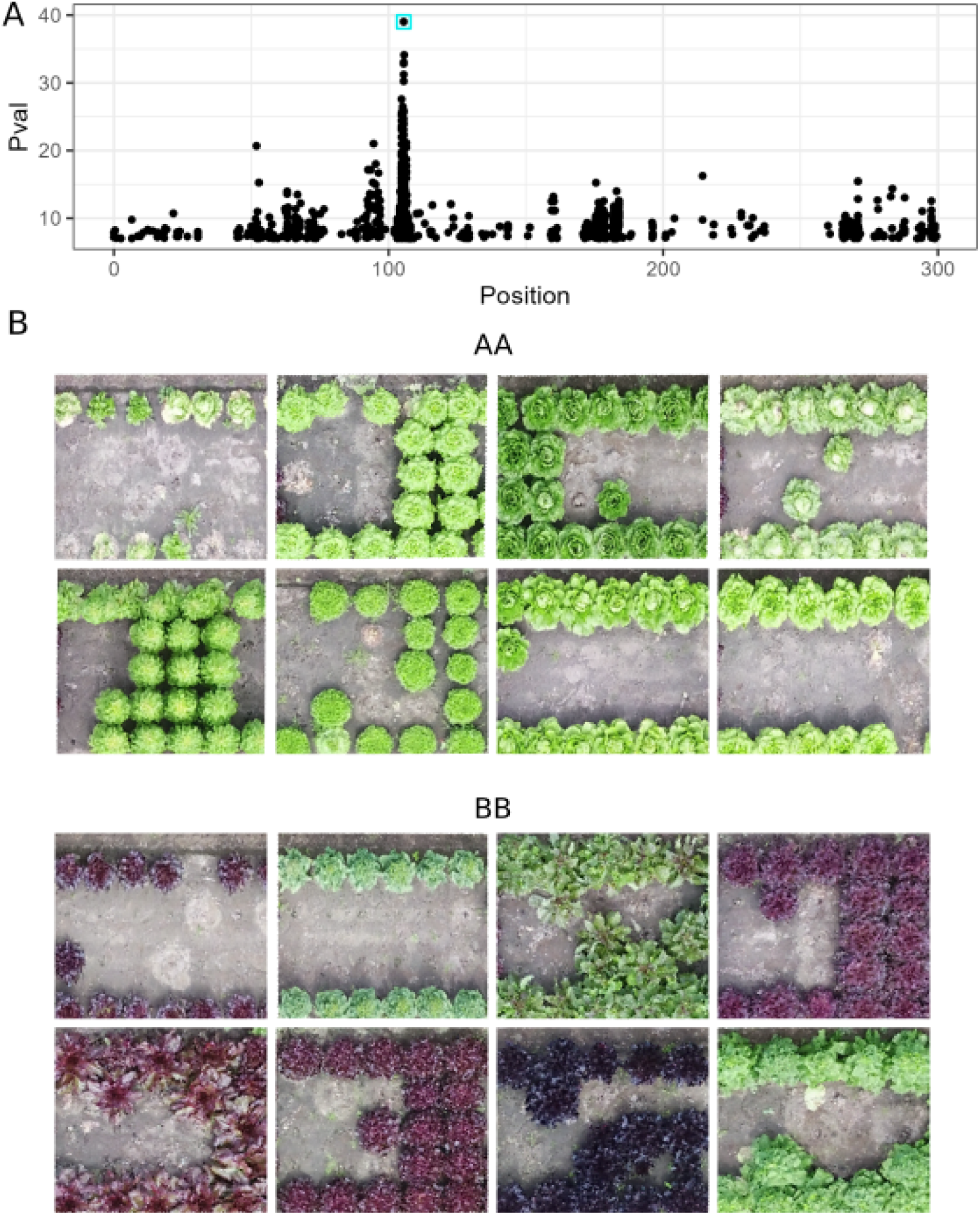
The traits causing the QTL on chromosome 8. . **A**) A zoom in on the combined results for all traits for chromosome 8 with the position in Mbp on the x-axis and the -log10(pvalue) on the y-axis. A highly significant SNP located at 105 Mbp is circled in teal. **B**) Accessions with contrasting alleles for the SNP circled in panel A. When looking at 8 random accessions homozygous for the reference allele we see green accessions. When examining all 8 accessions homozygous for the alternative allele 5 out of 8 accessions show the typic purple anthocyanin color, and the other three accessions are dark green.

**Supplementary Figure 8:**
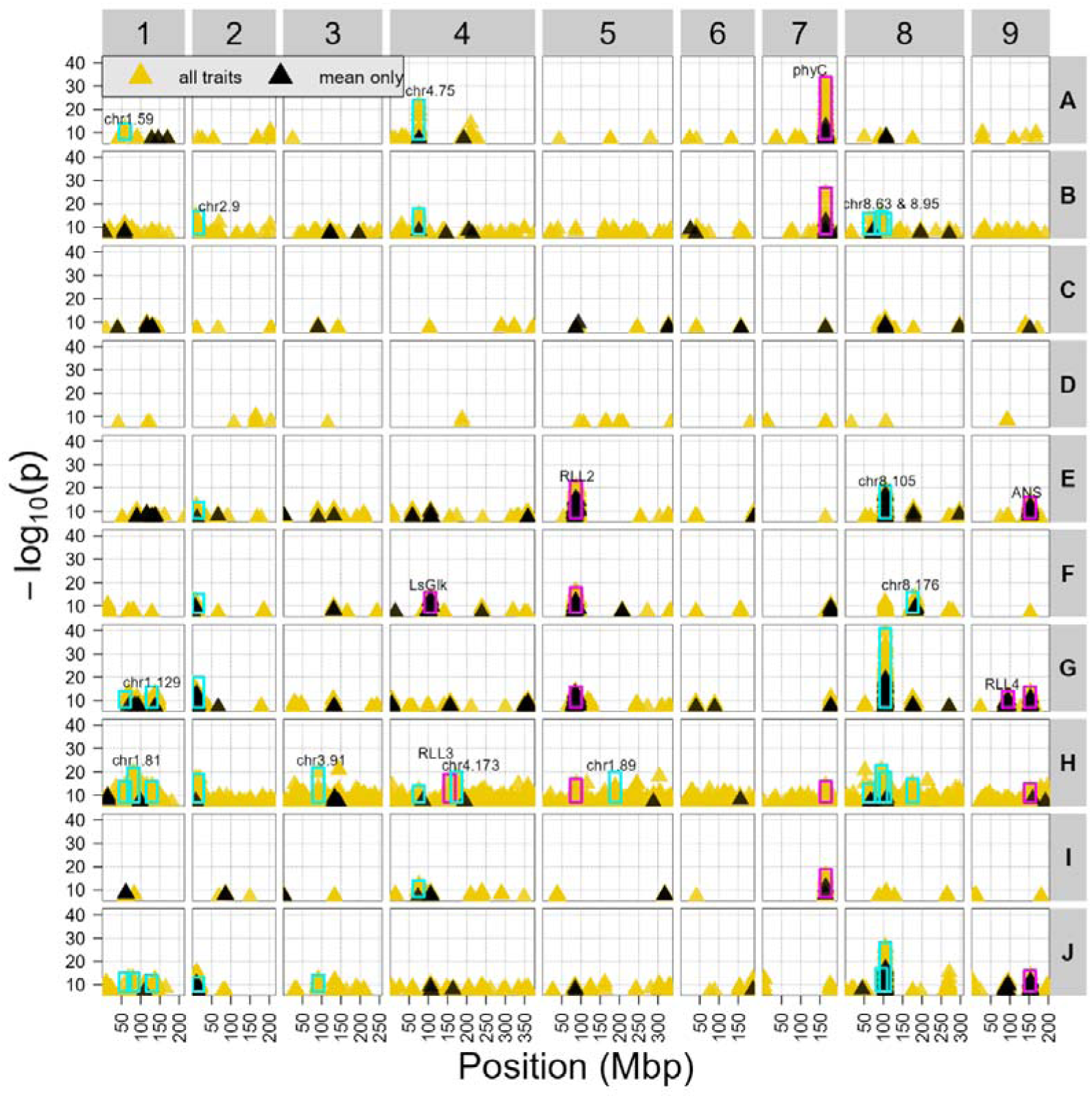
Comparison of using only the mean traits or using many extended descriptives. Every row represents a cluster of traits. The black dots represent a mean trait, while the yellow dots represent extended descriptives. QTLs that have been previously reported are annotated with a purple rectangle while (to our knowledge) new QTLs are shown with a teal rectangle.

**Supplementary Figure 9:**
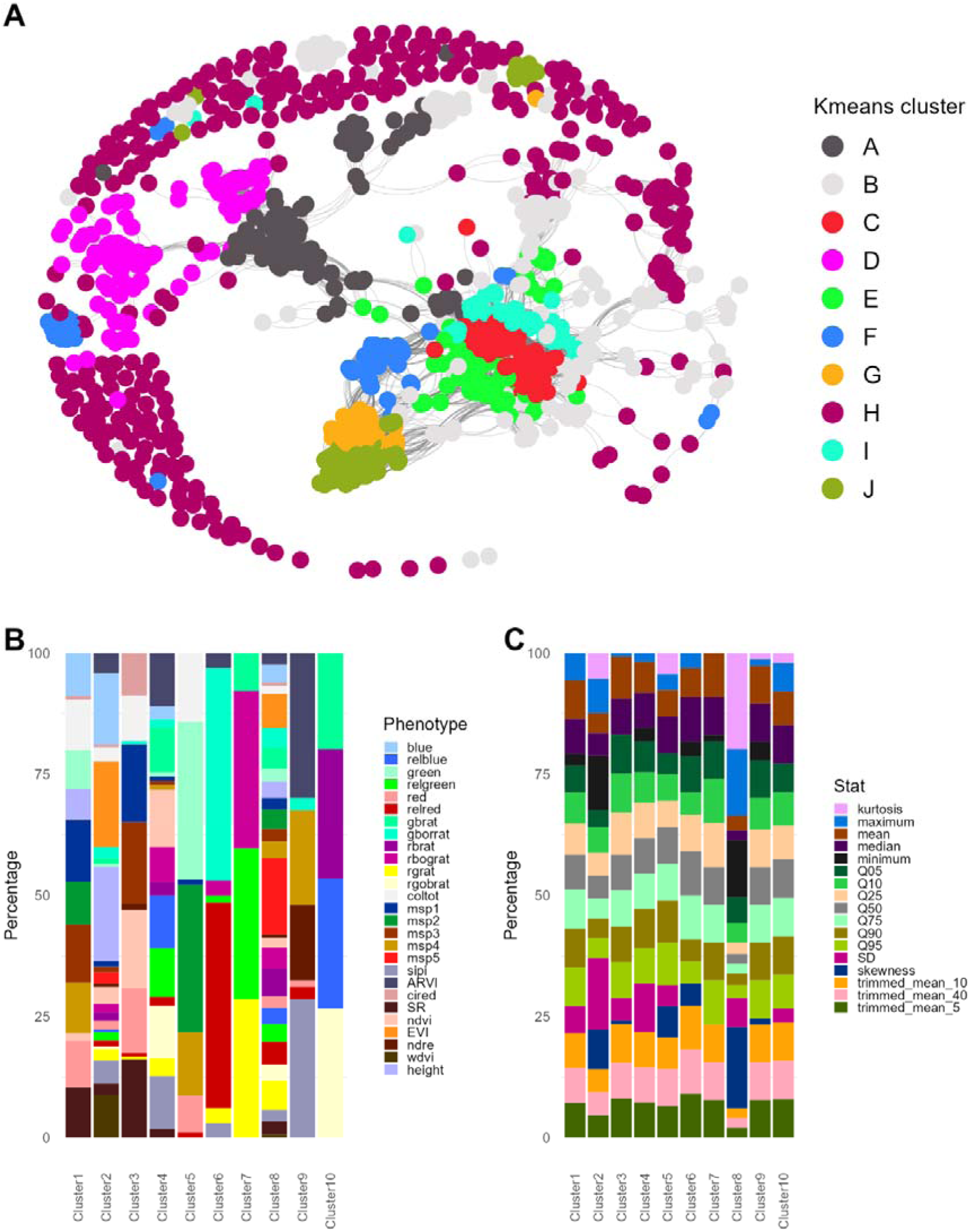
Details about the clustering on all traits, including extended descriptives. **A**) Network based on the trait*trait correlation matrix. Nodes represent traits, and edges represent correlations > …. Color shows the cluster assignment of k-means clustering (K = 10) on the correlation matrix of all traits. **B**) Distribution of phenotypes over clusters. **C**) Distribution of descriptives over clusters.

**Supplementary Figure 10:**
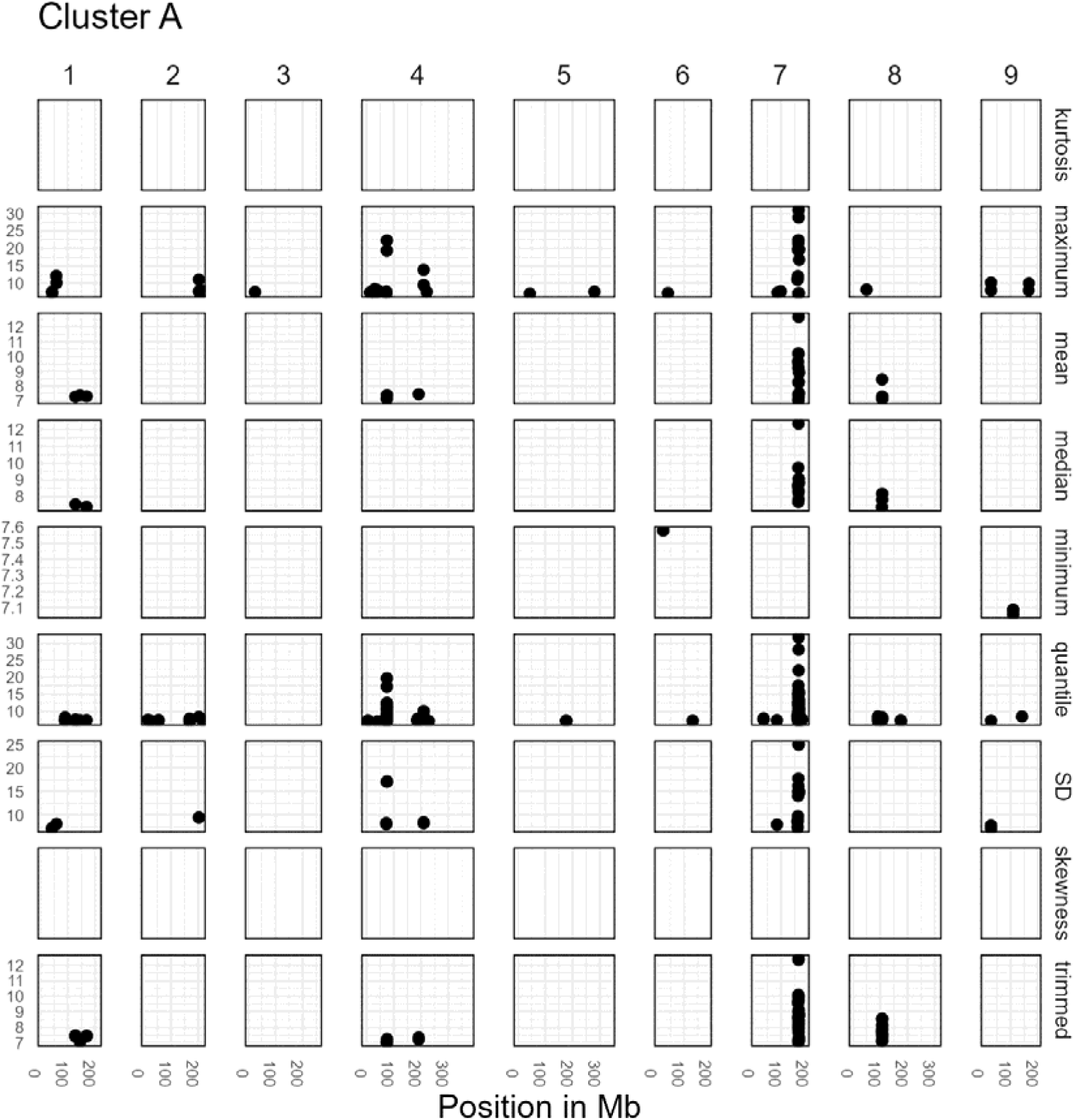
The QTLs found with traits from cluster A (in the clustering that includes the extended descriptives).

**Supplementary Figure 11:**
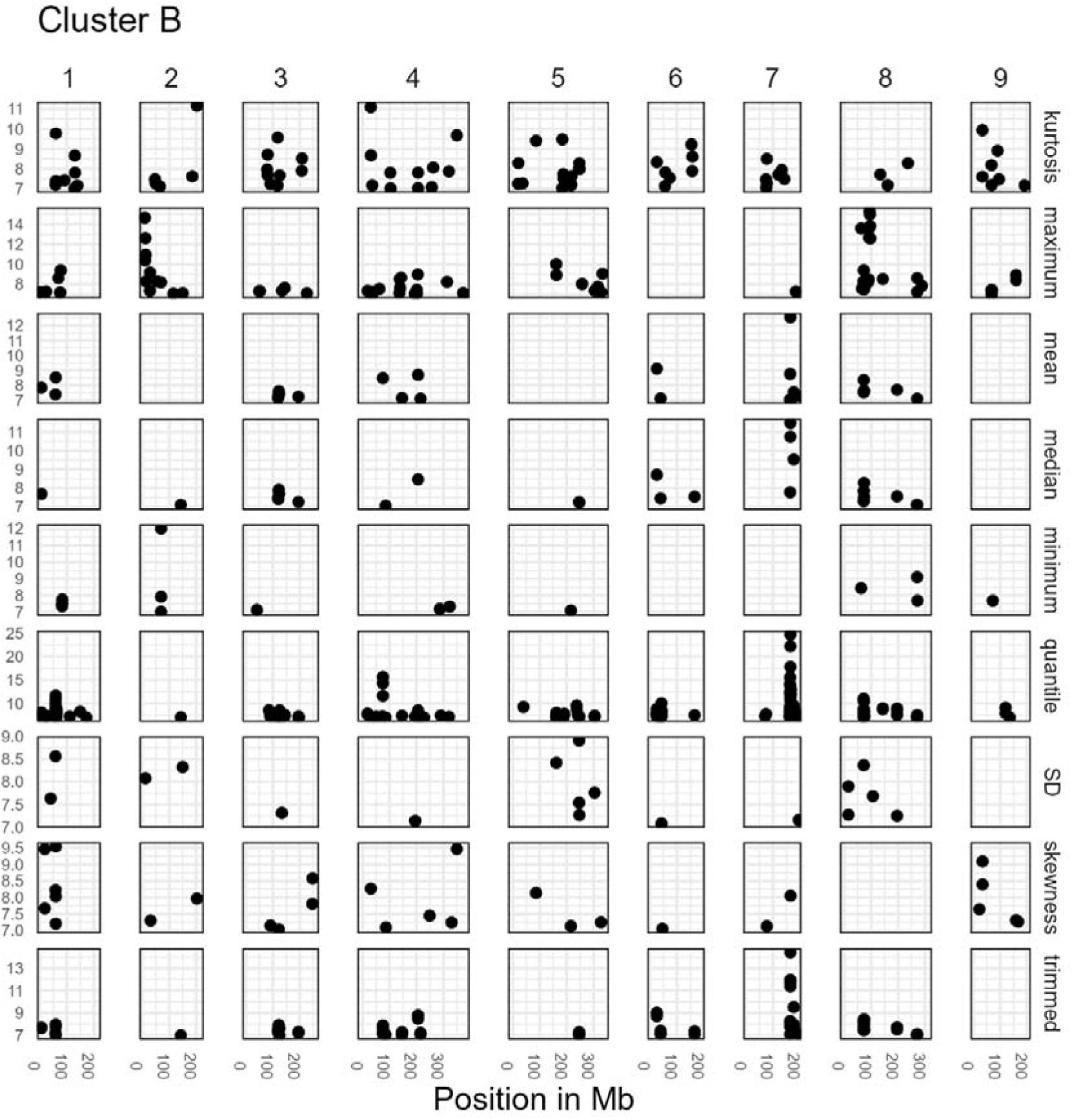
The QTLs found with traits from cluster B (in the clustering that includes the extended descriptives).

**Supplementary Figure 12:**
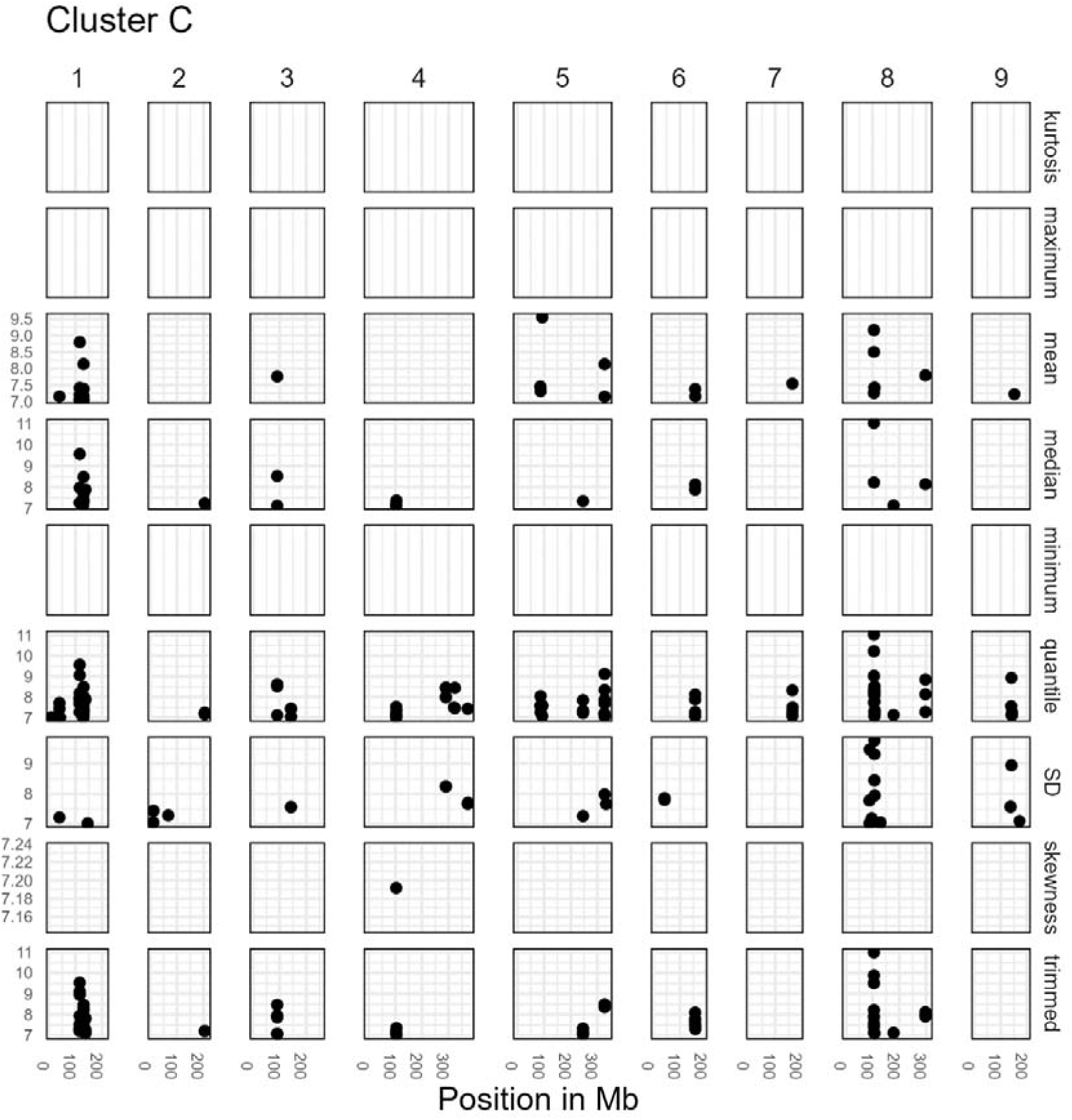
The QTLs found with traits from cluster C (in the clustering that includes the extended descriptives).

**Supplementary Figure 13:**
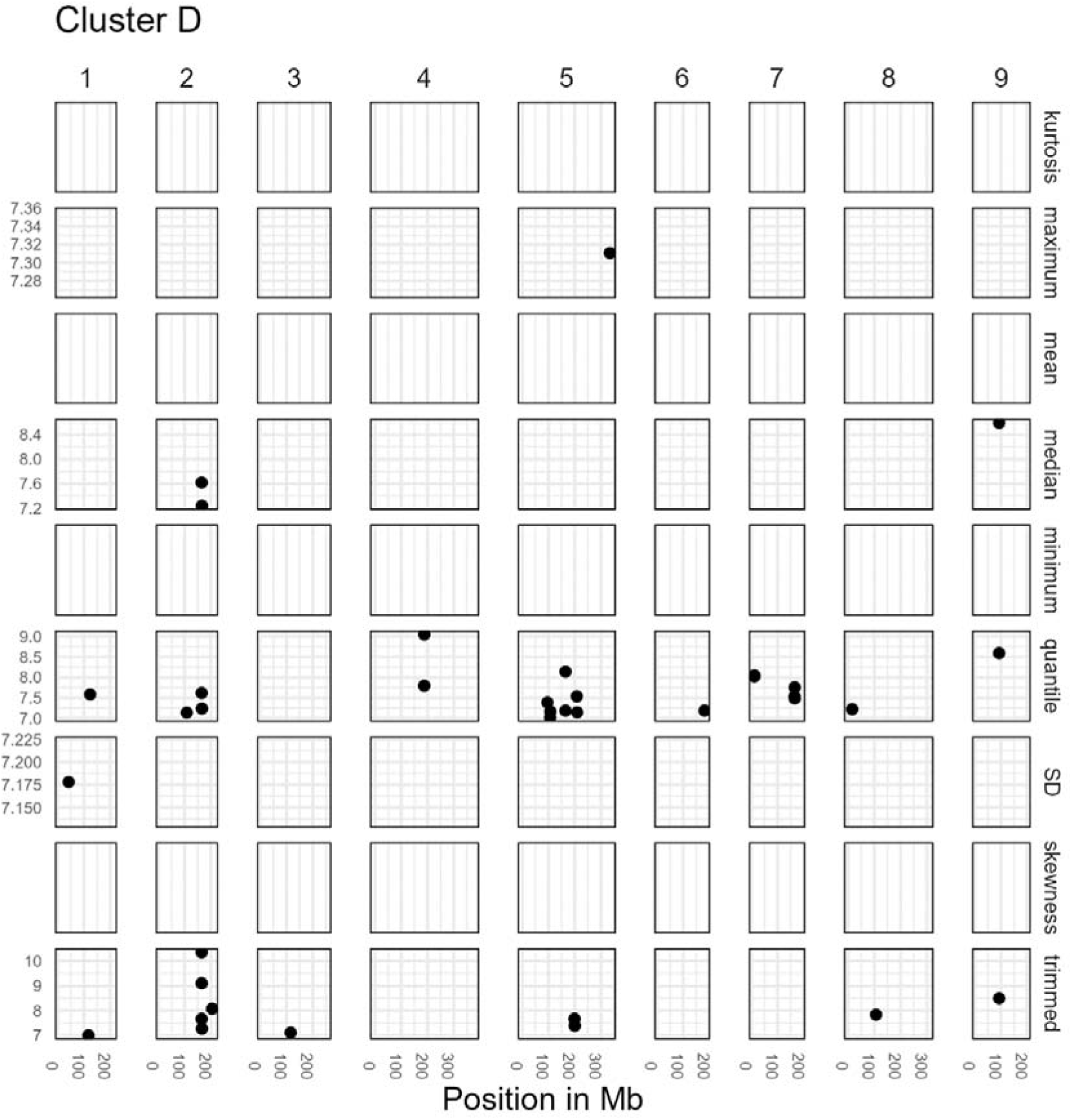
The QTLs found with traits from cluster D (in the clustering that includes the extended descriptives).

**Supplementary Figure 14:**
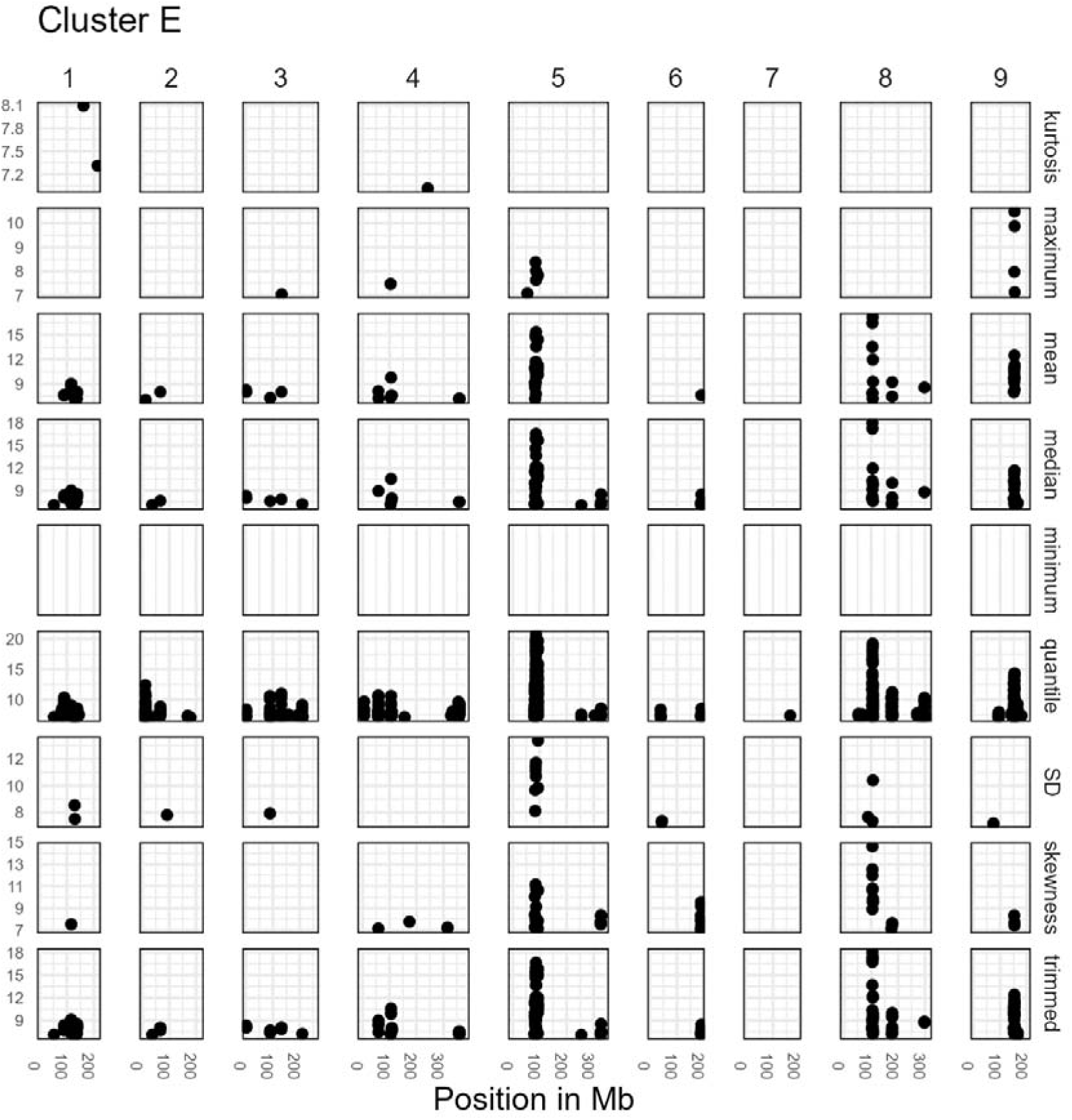
The QTLs found with traits from cluster E (in the clustering that includes the extended descriptives).

**Supplementary Figure 15:**
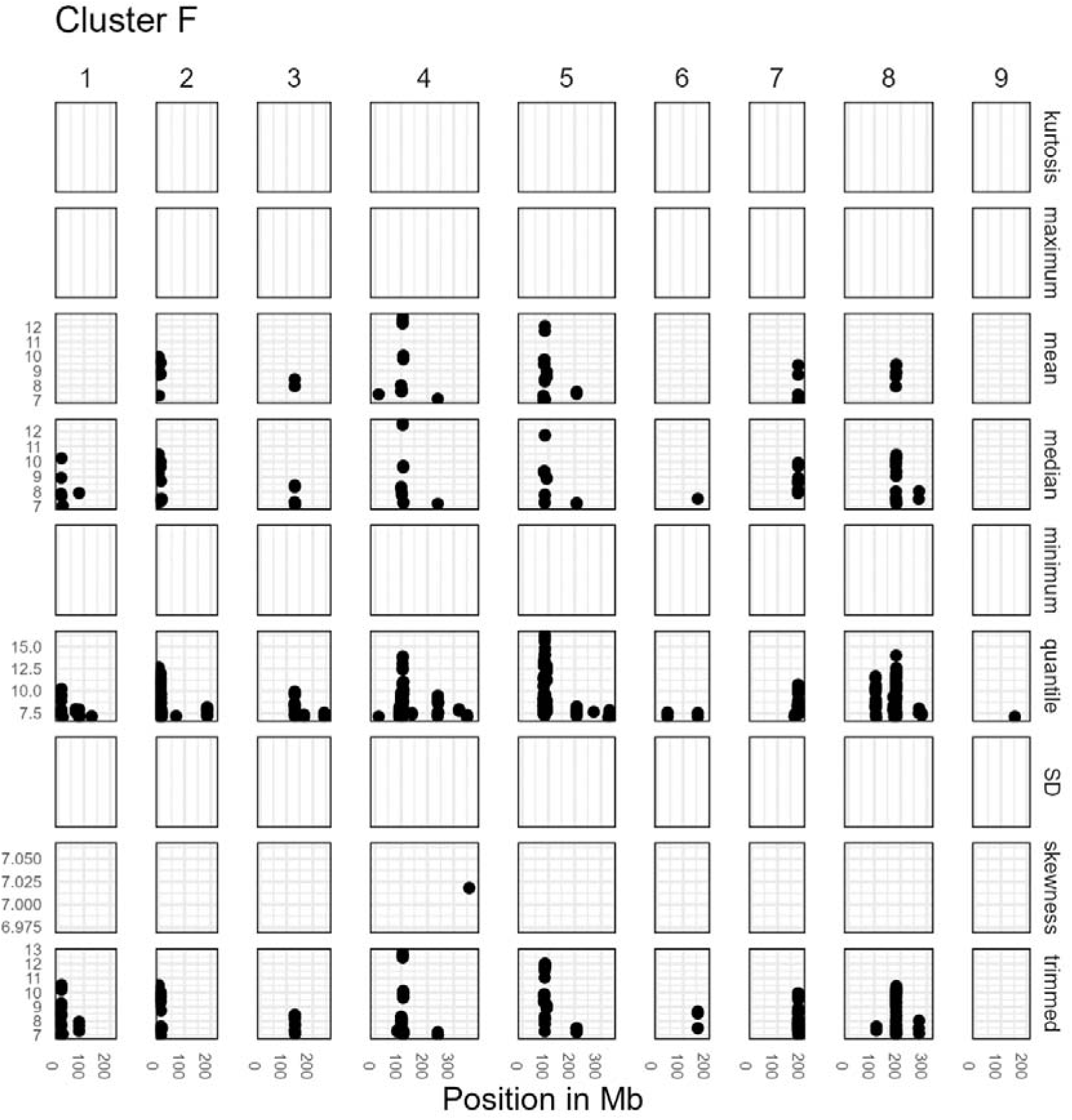
The QTLs found with traits from cluster F (in the clustering that includes the extended descriptives).

**Supplementary Figure 16:**
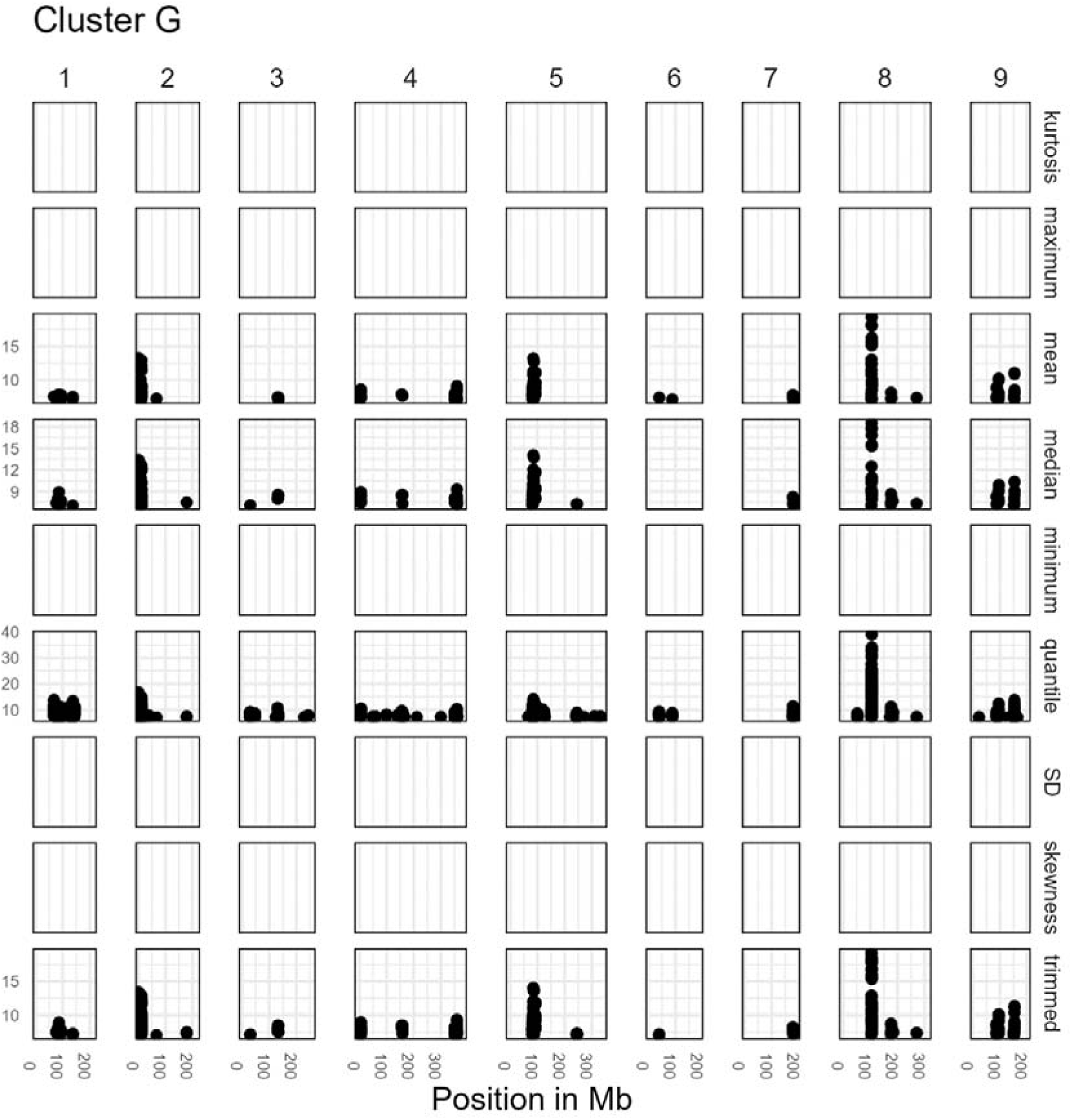
The QTLs found with traits from cluster G (in the clustering that includes the extended descriptives).

**Supplementary Figure 17:**
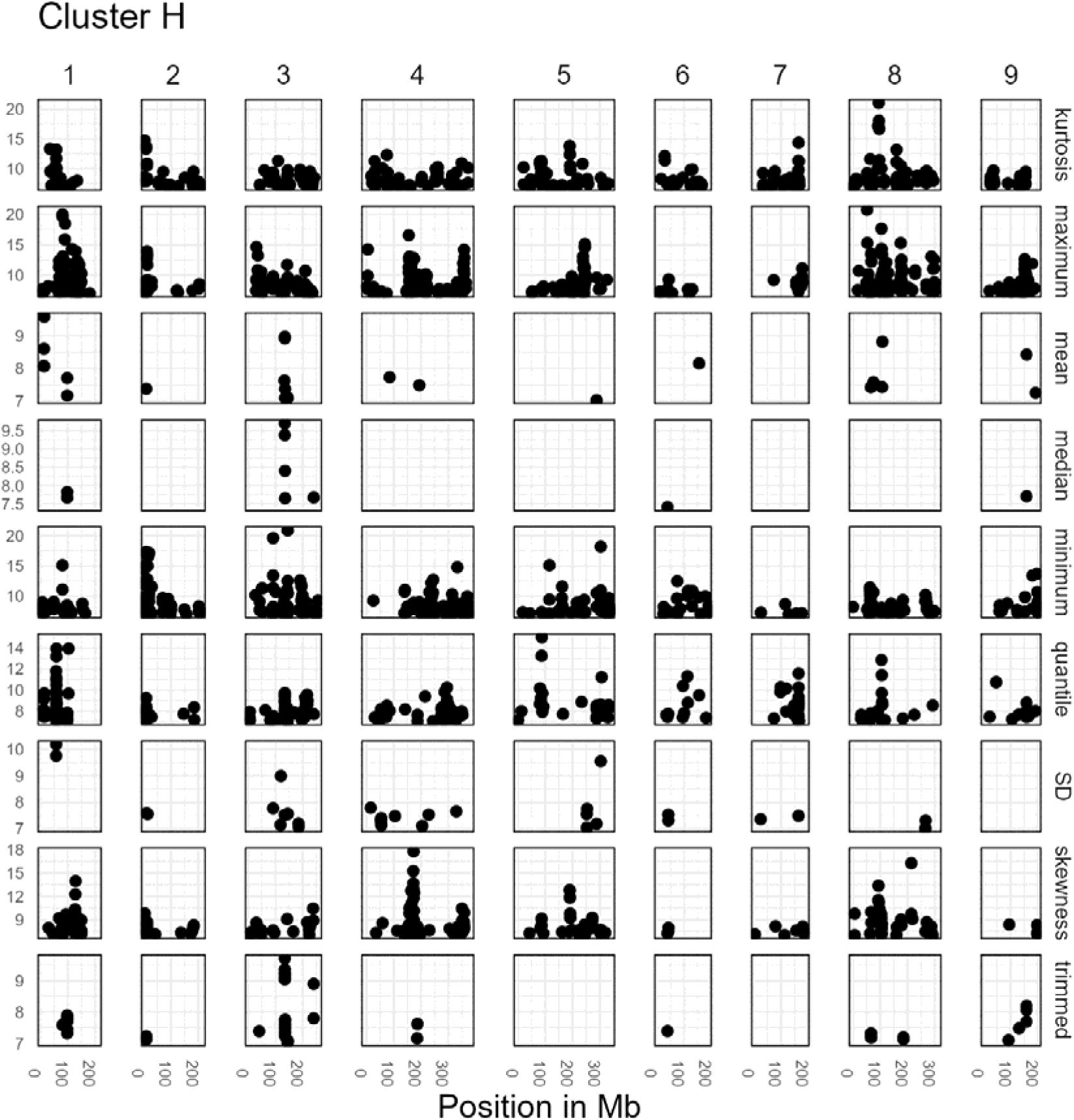
The QTLs found with traits from cluster H (in the clustering that includes the extended descriptives).

**Supplementary Figure 18:**
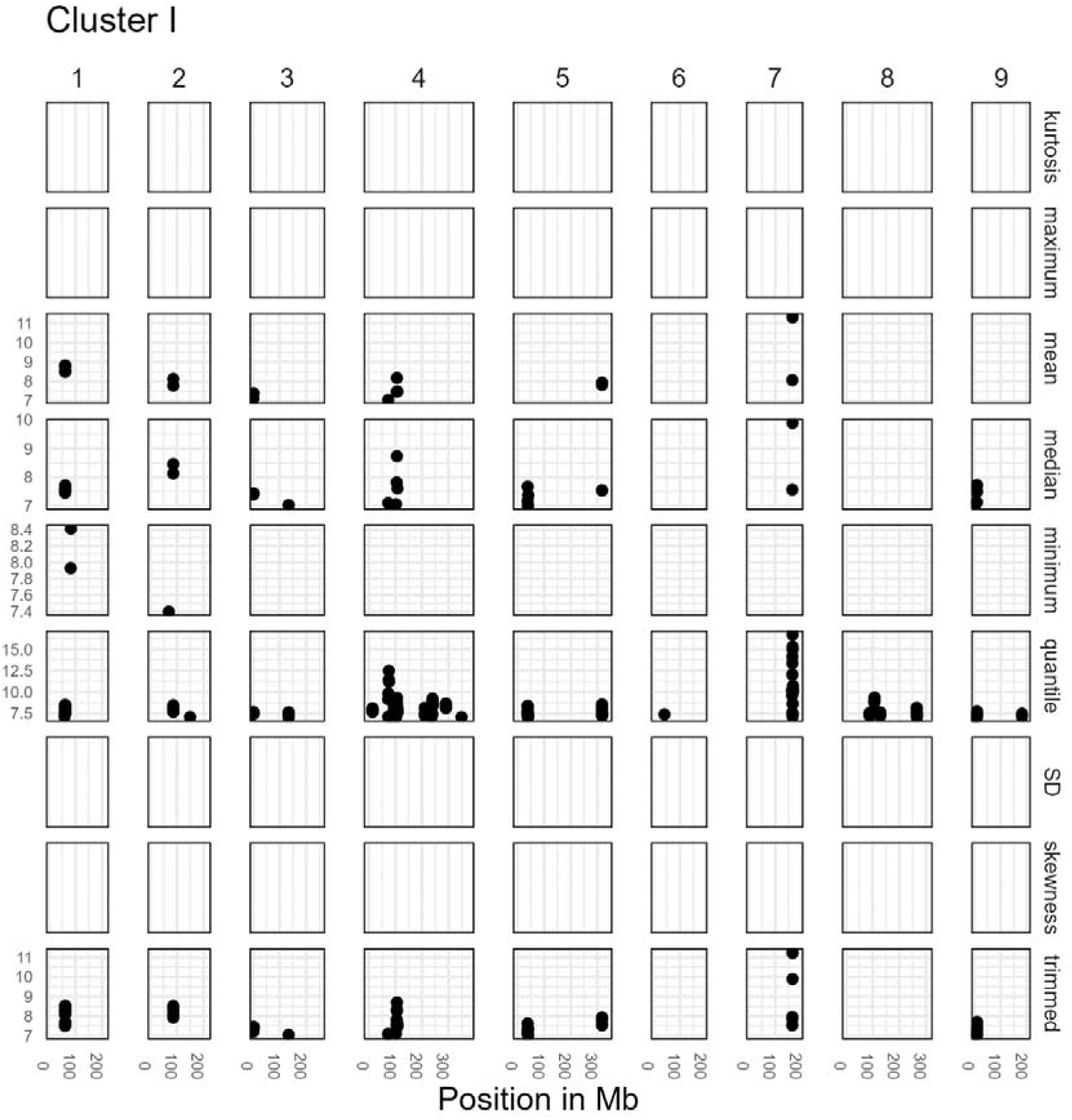
The QTLs found with traits from cluster I (in the clustering that includes the extended descriptives).

**Supplementary Figure 19:**
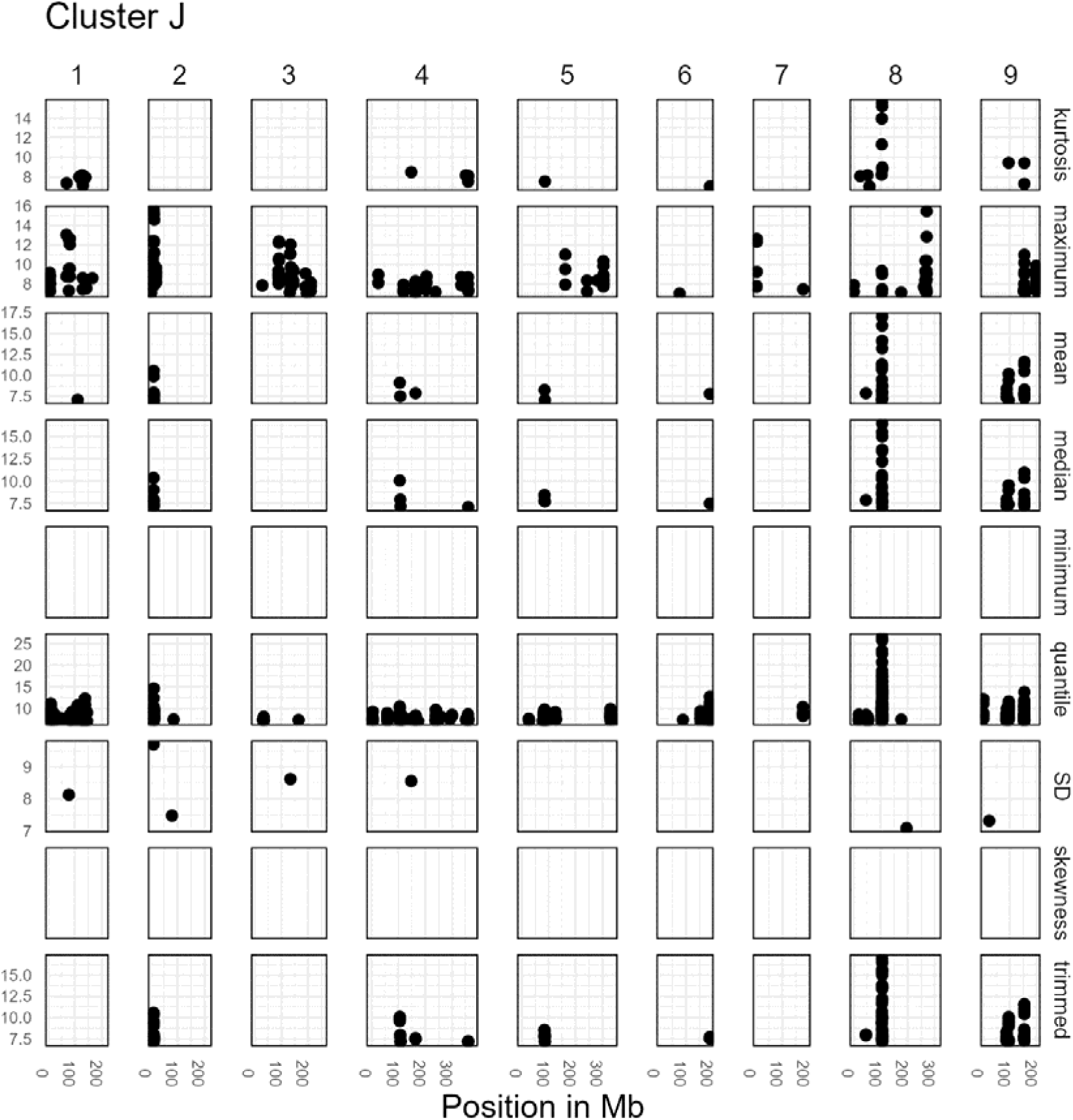
The QTLs found with traits from cluster J (in the clustering that includes the extended descriptives).

## References

Bac-Molenaar, J.A., Granier, C., Keurentjes, J.J.B. and Vreugdenhil, D. (2016) Genome-wide association mapping of time-dependent growth responses to moderate drought stress in Arabidopsis. Plant Cell Environment, 39, 88–102.

Briatte, F. (2023) Geometries to Plot Networks with “ggplot2” [R package ggnetwork version 0.5.12]. Available at: https://CRAN.R-project.org/package=ggnetwork [Accessed August 10, 2023].

Celesnik, H., Ali, G.S., Robison, F.M. and Reddy, A.S.N. (2013) Arabidopsis thaliana VOZ (Vascular plant One-Zinc finger) transcription factors are required for proper regulation of flowering time. Biology Open, 2, 424–431.

Chen, C., Huang, W., Hou, K. and Wu, W. (2019) Bolting, an important process in plant development, two types in plants. Journal of Plant Biology, 62, 161–169.

Chen, M., Ji, M., Wen, B., Liu, L., Li, S., Chen, X., Gao, D. and Li, L. (2016) GOLDEN 2-LIKE transcription factors of plants. Frontiers Plant Science, 7, 1509.

Cheng, D.M., Pogrebnyak, N., Kuhn, P., Krueger, C.G., Johnson, W.D. and Raskin, I. (2014) Development and Phytochemical Characterization of High Polyphenol Red Lettuce with Anti-Diabetic Properties. PLoS One, 9, e91571.

Cho, J.S., Nguyen, V.P., Jeon, H.W., et al.(2016) Overexpression of PtrMYB119, a R2R3-MYB transcription factor from Populus trichocarpa, promotes anthocyanin production in hybrid poplar. Tree Physiology, 36, 1162–1176.

Ding, G., Shen, L., Dai, J., et al. (2023) The Dissection of Nitrogen Response Traits Using Drone Phenotyping and Dynamic Phenotypic Analysis to Explore N Responsiveness and Associated Genetic Loci in Wheat. Plant Phenomics, 5, 0128.

DroneWorkers (2022) Home - DroneWerkers. Available at: https://www.dronewerkers.nl/english/ [Accessed November 14, 2023].

Eijnatten, A.L. van, Sterken, M.G., Kammenga, J.E., Nijveen, H. and Snoek, B.L. (2024) The effect of developmental variation on expression QTLs in a multi parental Caenorhabditis elegans population. G3 (Bethesda), 14, 2.

Falcioni, R., Gonçalves, J.V.F., Oliveira, K.M. de, Antunes, W.C. and Nanni, M.R.(2022) VIS-NIR-SWIR Hyperspectroscopy Combined with Data Mining and Machine Learning for Classification of Predicted Chemometrics of Green Lettuce. Remote Sensing *2022, Vol. 14, Page* 6330, 14, 6330.

Falcioni, R., Gonçalves, J.V.F., Oliveira, K.M. de, Oliveira, C.A. de, Demattê, J.A.M., Antunes, W.C. and Nanni, M.R. (2023) Enhancing Pigment Phenotyping and Classification in Lettuce through the Integration of Reflectance Spectroscopy and AI Algorithms. Plants, 12, 1333.

Han, R., Truco, M.J., Lavelle, D.O. and Michelmore, R.W. (2021) A Composite Analysis of Flowering Time Regulation in Lettuce. Frontiers in Plant Science, 12, 632708.

Han, R., Wong, A.J.Y., Tang, Z., Truco, M.J., Lavelle, D.O., Kozik, A., Jin, Y. and Michelmore, R.W. (2021) Drone phenotyping and machine learning enable discovery of loci regulating daily floral opening in lettuce. Journal of Experimental Botany, 72, 2979–2994.

Heimler, D., Isolani, L., Vignolini, P., Tombelli, S. and Romani, A. (2007) Polyphenol content and antioxidative activity in some species of freshly consumed salads. Journal of Agricultural and Food chemistry, 55, 1724–1729.

Huang, S., Tang, L., Hupy, J.P., Wang, Y. and Shao, G.(2021) A commentary review on the use of normalized difference vegetation index (NDVI) in the era of popular remote sensing. Journal of Forestry Research, 32, 1–6.

Kureel, N., Sarup, J., Matin, S., Goswami, S. and Kureel, K. (2022) Modelling vegetation health and stress using hypersepctral remote sensing data. Modeling Earth Systems and Environment 8, 733–748.

Laskowski, M.J., Dreher, K.A., Gehring, M.A., Abel, S., Gensler, A.L. and Sussex, I.M. (2002) FQR1, a Novel Primary Auxin-Response Gene, Encodes a Flavin Mononucleotide-Binding Quinone Reductase. Plant Physiology, 128, 578–590.

Leijten, W., Koes, R., Roobeek, I. and Frugis, G. (2018) Translating Flowering Time from Arabidopsis thaliana to Brassicaceae and Asteraceae Crop Species. Plants *2018, Vol. 7, Page 111*, 7, 111.

Mampholo, B.M., Maboko, M.M., Soundy, P. and Sivakumar, D. (2016) Phytochemicals and Overall Quality of Leafy Lettuce (Lactuca sativa L.) Varieties Grown in Closed Hydroponic System. Journal of Food Quality, 39, 805–815.

Poplin, R., Ruano-Rubio, V., DePristo, M.A., et al. (2018) Scaling accurate genetic variant discovery to tens of thousands of samples. bioRxiv, 201178.

R Core Team (2023) R: A Language and Environment for Statistical Computing.

Rosental, L., Still, D.W., You, Y., Hayes, R.J. and Simko, I.(2021) Mapping and identification of genetic loci affecting earliness of bolting and flowering in lettuce. Theoretical and Applied Genetics *2021 134:10*, 134, 3319–3337.

Shi, M., Gu, J., Wu, H., et al. (2022) Phytochemicals, Nutrition, Metabolism, Bioavailability, and Health Benefits in Lettuce—A Comprehensive Review. Antioxidants *2022, Vol. 11, Page 1158*, 11, 1158.

Spindel, J.E., Dahlberg, J., Colgan, M., Hollingsworth, J., Sievert, J., Staggenborg, S.H., Hutmacher, R., Jansson, C. and Vogel, J.P. (2018) Association mapping by aerial drone reveals 213 genetic associations for Sorghum bicolor biomass traits under drought. BMC Genomics, 19, 1–18.

Su, W., Tao, R., Liu, W., et al.(2020) Characterization of four polymorphic genes controlling red leaf colour in lettuce that have undergone disruptive selection since domestication. Plant Biotechnology Journal, 18, 479–490.

Vasseur, F., Bontpart, T., Dauzat, M., Granier, C. and Vile, D.(2014) Multivariate genetic analysis of plant responses to water deficit and high temperature revealed contrasting adaptive strategies. Journal of experimental botany, 65, 6457–6469.

Watanabe, K., Guo, W., Arai, K., et al. (2017) High-throughput phenotyping of sorghum plant height using an unmanned aerial vehicle and its application to genomic prediction modeling. Frontiers in Plant Science, 8, 254051.

Wei, T., Treuren, R. van, Liu, Xinjiang, et al. (2021) Whole-genome resequencing of 445 Lactuca accessions reveals the domestication history of cultivated lettuce. Nature Genetics *2021 53:5*, 53, 752–760.

Wilmouth, R.C., Turnbull, J.J., Welford, R.W.D., Clifton, I.J., Prescott, A.G. and Schofield, C.J. (2002) Structure and Mechanism of Anthocyanidin Synthase from Arabidopsis thaliana and have long-established biomedicinal properties, including inhibition of cell proliferation and antimutagenic, antimicrobial, antiinflammatory, and antihypertensive. Structure, 10, 93–103.

Workum, D.-J.M. Van, Mehrem, S.L., Basten, et al. (2024a) Lactuca super-pangenome reduces bias towards reference genes in lettuce research. bioRxiv, 2024.06.20.599299.

Workum, D.-J.M. Van, Ridder, D. de, Schranz, M.E. and Smit, S. (2024b) Dataset: Structural, functional and evolutionary characterisation of genes in Lactuca sp. reference genomes in the context of eudicots (dataset). datacite.

Xiao, Q., Bai, X., Zhang, C. and He, Y. (2022) Advanced high-throughput plant phenotyping techniques for genome-wide association studies: A review. Journal of advanced Research, 35, 215–230.

Yamaguchi, A., Wu, M.F., Yang, L., Wu, G., Poethig, R.S. and Wagner, D.(2009) The MicroRNA-Regulated SBP-Box Transcription Factor SPL3 Is a Direct Upstream Activator of LEAFY, FRUITFULL, and APETALA1. Developmental Cel,l 17, 268–278.

Yang, X., Gil, M.I., Yang, Q. and Tomás-Barberán, F.A. (2022) Bioactive compounds in lettuce: Highlighting the benefits to human health and impacts of preharvest and postharvest practices. Comprehensive Reviews in Food Science and Food Safety, 21, 4–45.

Ye, Y., Wang, P., Zhang, M., et al.(2023) UAV-based time-series phenotyping reveals the genetic basis of plant height in upland cotton. The Plant Journal, 115, 937– 951.

Zhang, L., Qian, J., Han, Y., Jia, Y., Kuang, H. and Chen, J. (2022) Alternative splicing triggered by the insertion of a CACTA transposon attenuates LsGLK and leads to the development of pale-green leaves in lettuce. The Plant Journal, 109, 182–195.

Zhang, L., Su, W., Tao, R., et al. (2017) RNA sequencing provides insights into the evolution of lettuce and the regulation of flavonoid biosynthesis. Nature Communications *2017 8:1*, 8, 1–12.

Zhang, Z., Treuren, R. van, Yang, T., Hu, Y., Zhou, W., Liu, H. and Wei, T.(2023) A comprehensive lettuce variation map reveals the impact of structural variations in agronomic traits. BMC Genomics, 24.

Ziyatdinov, A., Vázquez-Santiago, M., Brunel, H., Martinez-Perez, A., Aschard, H. and Soria, J.M. (2018) lme4qtl: Linear mixed models with flexible covariance structure for genetic studies of related individuals. BMC Bioinformatics, 19.

